# Metabolic dyshomeostasis induced by SARS-CoV-2 structural proteins reveals immunological insights into viral olfactory interactions

**DOI:** 10.1101/2022.02.01.478724

**Authors:** Mercedes Lachén-Montes, Naroa Mendizuri, Karina Ausín, Miriam Echaide, Ester Blanco, Luisa Chocarro, María de Toro, David Escors, Joaquín Fernández-Irigoyen, Grazyna Kochan, Enrique Santamaría

## Abstract

One of the most common symptoms in COVID-19 is a sudden loss of smell. SARS-CoV-2 has been detected in the olfactory bulb (OB) from animal models and sporadically in COVID-19 patients. To decipher the specific role over the SARS-CoV-2 proteome at olfactory level, we characterized the in-depth molecular imbalance induced by the expression of GFP-tagged SARS-CoV-2 structural proteins (M, N, E, S) on mouse OB cells. Transcriptomic and proteomic trajectories uncovered a widespread metabolic remodeling commonly converging in extracellular matrix organization, lipid metabolism and signaling by receptor tyrosine kinases. The molecular singularities and specific interactome expression modules were also characterized for each viral structural factor. The intracellular molecular imbalance induced by each SARS-CoV-2 structural protein was accompanied by differential activation dynamics in survival and immunological routes in parallel with a differentiated secretion profile of chemokines in OB cells. Machine learning through a proteotranscriptomic data integration uncovered TGF-beta signaling as a confluent activation node by the SARS-CoV-2 structural proteome. Taken together, these data provide important avenues for understanding the multifunctional immunomodulatory properties of SARS-CoV-2 M, N, S and E proteins beyond their intrinsic role in virion formation, deciphering mechanistic clues to the olfactory inflammation observed in COVID-19 patients.

## 1. Introduction

The coronavirus disease 2019 (COVID-19) is an ongoing viral pandemic caused by the rapid spread of severe acute respiratory syndrome coronavirus 2 (SARS-CoV-2) (Li et al., 2020). It is well known that infection is transmitted by respiratory droplets with an incubation period of approximately 5 days (Stadnytskyi et al., 2020), however, there are still many gaps regarding the molecular mechanisms disrupted during the infection and the immune system response triggered against this pathogen. The approximately 30,000-nucleotide SARS-CoV-2 RNA genome encodes for 14 major open-reading frames (ORFs). The 4 structural proteins (S (spike), membrane (M), envelope (E), and nucleocapside (N)) are expressed from subgenomic RNAs transcribed from the RNA genome. In addition, SARS-CoV-2 expresses another 14 non-structural proteins and 8 accessory proteins which participate in RNA replication-transcription, and pathogenesis (Naqvi et al., 2020). The clinical presentation of COVID-19 varies from asymptomatic, mild to moderate self-limiting disease in the majority of cases (Huang et al., 2020). However, fatal consequence can occur in patients with comorbidities (Guzik et al., 2020;Yang et al., 2020). SARS-CoV-2 infection in human patients include fever and respiratory symptoms, such as fever, dry cough, dyspnea, and pneumonia (Huang et al., 2020). In addition, accumulating evidence has demonstrated the appearance of other symptoms of clinical importance, such as neurological affectation and olfactory and gustatory dysfunctions, among others (Bagheri et al., 2020;Beltran-Corbellini et al., 2020;Giacomelli et al., 2020;Menni et al., 2020;Parma et al., 2020;Gerkin et al., 2021). Importantly from a diagnostic point of view, these changes usually occur earlier than other symptoms, suggesting a potential specific predictive value (Haehner et al., 2020;Pierron et al., 2020).

Although the molecular mechanisms of olfactory and neurological disorders in SARS-CoV-2-infected patients are not fully understood, there is growing evidence that the olfactory loss observed in COVID-19 occurs through the infection of supporting cells in the olfactory system (Brann et al., 2020). The key pathological mechanism of action for SARS-CoV-2 to invade the host cell involves the S protein attachment to the human angiotensin-converting enzyme 2 (ACE2) cell surface protein. Then, the virus uptake is facilitated by the action of the TMPRSS2 protease (Shang et al., 2020). Importantly, sustentacular cells highly express both ACE2 and TMPRSS2 proteins, while little expression is found in olfactory neurons (Brann et al., 2020). Therefore, these sustentacular cells might be the main targets for SARS-CoV-2 infection, leaving the neurons vulnerable and depleted of nutrients. However, other studies which focused on the olfactory epithelium from SARS-CoV-2-infected patients with acute loss of smell, point out that olfactory sensory neurons, support cells, and immune cells are major sites of SARS-CoV-2 infection (de Melo et al., 2021). Certain coronaviruses have the ability to pass from the OE through the cribriform plate to infect the OB (Durrant et al., 2016;Dube et al., 2018). In fact, several reports have detected evidences of SARS-CoV-2 at the OB (Lou et al., 2021). However, the neuroinvasive potential of SARS-CoV-2 and future neurological vulnerabilities remain controversial (Cooper et al., 2020;Xydakis et al., 2021).

Omics technologies have allowed to increase our knowledge about the molecular imbalance triggered by the SARS-CoV-2 in multiple human organs as well as in-vitro systems (Lachen-Montes et al., 2020;Nie et al., 2021;Saccon et al., 2021;Stukalov et al., 2021). However, there are still major gaps in the pathways leading to loss of smell caused by SARS-CoV-2, in addition to the molecular consequences from SARS-CoV-2 structural proteins at the olfactory level. Bearing in mind that the olfactory loss is considered an early predictor of COVID-19, we consider that the methodologies simulating SARS-CoV-2 infection in olfactory cells in combination with -omics strategies is a straightforward approach to elucidate the host and viral mechanisms involved in SARS-CoV-2-induced anosmia. These approaches can also identify potential therapeutic targets for non-invasive intranasal treatments. In this study, we have characterized the molecular dyshomeostasis induced by each SARS-CoV-2 structural protein (S, N, M, E) in murine OBC1 cells as an olfactory in-vitro system. The effects induced by the expression of GFP-tagged SARS-CoV-2 structural proteins uncovered a tangled crosstalk between multiple biofunctions and pathways as well as an immunological response differentially modulated by each SARS-CoV-2 structural protein. Our results also identified novel viral-host protein interactions partially shared by the structural components of the SARS-CoV-2 proteome.

## 2. Materials and Methods

### 2.1 Materials

The following reagents and materials were used. Anti-p38 MAPK (ref. 9212), anti-phospho-p38 MAPK (T180/Y182) (ref. 9211), anti-Akt (ref. 4685), anti-phospho-Akt (S473) (ref. 4060), anti-PKA C-alpha (ref. 4782), anti-phospho-PKA C (T197) (ref. 5661), anti-SEK1 (ref. 9152), anti-phospho-SEK1 (S257/T261) (Ref. 9156), anti-MEK1/2 (ref. 9126), anti-phospho-MEK1/2 (S217/221) (ref. 9154), anti-phospho GSK3 α/β (S21) (ref. 9331), anti-GSK3 α/β (ref. 5676), anti-Phb1 (ref. 2426), and anti-Phb2 (ref. 14085) anti-PDK1 (ref. 3062), anti-phospho-PDK1 (S241) (ref. 3061), anti-phospho-PKC pan (T514) (ref. 9379), anti-MEK1/2 (ref. 9126), anti-phospho-MEK1/2 (S217/221) (ref. 9154), anti-phospho CAMKII (Y86) (ref.12716) and anti-CAMKII (ref.11945) were purchased from Cell Signaling Technology. Anti-PKC-pan was from Sigma Aldrich (ref. SAB4502356). Electrophoresis reagents were purchased from Bio-rad and trypsin from Promega.

### 2.2 Generation of M, E, N, S and GFP fusion proteins

Coding sequences for M, E, N and S proteins were retrieved from the SARS-CoV-2 reference genome (https://www.ncbi.nlm.nih.gov/nuccore/1798174254). Coding sequences were further codon-optimized for their expression in human cells, and recombinant synthetic genes fused to the GFP-coding sequence at the 3-termini were ordered from GeneArt (ThermoFisher). Genes were further cloned into pDUAL-PuroR lentivectors (Escors et al., 2008;Gato-Canas et al., 2017) under the transcriptional control of the SFFV promoter. pDUAL-PuroR lentivectors express puromycin resistance under the transcriptional control of the human ubiquitin promoter.

### 2.3 Cell culture and generation of transfected cell lines

Immortalized murine olfactory bulb OBC1 cells (ABM. T0235) were cultured in DMEM (Gibco) supplemented with 10% FBS (Merck Millipore) and 1% penicillin/streptomycin (ABM) and grown in a 5% CO_2_ humidified atmosphere at 37°C. For transfection, OBC1 cells were seeded in 6-well plates at a confluency of 80%. Twenty microliters of Fugene HD (Promega) were incubated separately in 50 ul of DMEM without FBS and then mixed with three micrograms of M-GFP, E-GFP, N-GFP, S-GFP and control GFP plasmid DNA, followed by 15 min incubation at room temperature. The cells were transfected in a 3ml total volume for 72 hours. After three days, the culture medium was replaced with DMEM supplemented with 1ug/ml puromycin (Thermo). Transfection efficiency was assessed using flow cytometry and to achieve a more specific population, cell sorting based on GFP expression was performed among all experimental conditions.

### 2.4 RNA sequencing (RNA-seq) and data analysis

Briefly, total RNA was extracted and purified using a RNeasy Mini Kit (Qiagen, Hilden, Germany) following manufacturer’s instructions. Sequencing libraries were prepared by following the Illumina Stranded Total RNA Prep with Ribo-Zero Plus (Illumina Inc., San Diego, CA) from 100 ng of total RNA, that has been depleted by following the instructions. All libraries were run in a HiSeq1500 PE100 lane in Rapid mode, pooled in equimolar amounts to a 10nm final concentration. The library concentration was measured by Qubit 3.0 (Invitrogen) and library size ensured by capilar electrophoresis in Fragment Analyzer (AATI). The quality of the RNAseq results was initially assessed using FastQC v0.11.9 (http://www.bioinformatics.babraham.ac.uk/projects/fastqc/) and MultiQC v1.9 (http://multiqc.info/). The raw reads were trimmed, filtered for those with a Phred quality score of at least 25 and all adapters were removed with TrimGalore v0.5.0 (https://www.bioinformatics.brabraham.ac.uk/projects/trim_galore/). Trimmed reads were analyzed with SortMeRNA v2.1 software (https://bioinfo.lifl.fr/RNA/sortmerna/) (Kopylova et al., 2012) to delete the 18S and 28S rRNA to eliminate the rRNA residues that could remain undepleted by the chemical treatment in the library preparation. Clean reads were aligned versus the Mus Musculus reference genome (release GRCm38.p6/GCA_000001635.8, ftp://ftp.ensembl.org) using HISAT2 v2.2.1 (https://daehwankimlab.github.io/hisat2/) (Kim et al., 2019) with default parameters. Resulting alignment files were quality assessed with Qualimap2 (http://qualimap.bioinfo.cipf.es) (Okonechnikov et al., 2016) and sorted and indexed with Samtools software (Li et al., 2009). After taking a read count on gene features with the FeatureCounts tool (http://subread.sourceforge.net) (Liao et al., 2014), quantitative differential expression analysis between conditions was performed by DESeq2 (Love et al., 2014), implemented as R Bioconductor package, performing read-count normalization by following a negative binomial distribution model. In order to automate this process and facilitate all group combination analysis, the SARTools pipeline (Varet et al., 2016) was used. All resultant data was obtained as HTML files and CSV tables, including density count distribution analysis, pairwise scatter plots, cluster dendrograms, Principal Component Analysis (PCoA) plots, size factor estimations, dispersion plots and MA and Volcano plots. The resulting CSV file, including raw counts, normalized counts, Fold-Change estimation and dispersion data was annotated with additional data from the Biomart database (https://www.ensembl.org/biomart/martview/346d6d487e88676fd509a1b9a642edb2). In order to control the False Discovery Rate (FDR), the p-values were amended by Benjamini-Hochberg (BH) multiple testing corrections. Those features showing corrected p-values below the 0.05 threshold and FoldChange values >1.5 or <0.5 were considered up- or down-regulated genes, respectively.

### 2.5 Proteomics and data analysis

#### 2.5.1 Protein extraction

Culture medium was removed, and cells were washed with 1X cold PBS. Then, pellet was resuspended in lysis buffer containing 7M urea, 2M tiourea and 50Mm DTT and incubated in ice for 30 min, vortexing each 10 min. After a sonication step, the lysate was centrifuged for 20 minutes at 20000xg at 15°C. The supernantant was then added to a new Eppendorf and protein concentration was assessed using the Bradford assay (Biorad).

#### 2.5.2 SWATH-mass spectrometry proteomics: MS/MS library generation

As an input for generating the SWATH-MS assay library, a pool of 15 samples (2.5μg/sample) derived each cell replicate was used. Protein extracts (30 μg) were diluted in Laemmli sample buffer and loaded into a 1.5 mm thick polyacrylamide gel with a 4% stacking gel casted over a 12.5% resolving gel. Total gel was stained with Coomassie Brilliant Blue and 12 equals slides from the pooled sample was excised from the gel and transferred into 1.5 mL Eppendorf tubes. Protein enzymatic cleavage was carried out with trypsin (Promega; 1:20, w/w) at 37 °C for 16 h. Peptide mixture was dried in a speed vacuum for 20 min. Purification and concentration of peptides was performed using C18 Zip Tip Solid Phase Extraction (Millipore). Peptides recovered from in-gel digestion processing were reconstituted into a final concentration of 0.5μg/μL of 2% ACN, 0.5% FA, 97.5% MilliQ-water prior to mass spectrometric analysis. MS/MS datasets for spectral library generation were acquired on a Triple TOF 5600+ mass spectrometer (Sciex, Canada) interfaced to an Eksigent nanoLC ultra 2D pump system (SCIEX, Canada) fitted with a 75 μm ID column (Thermo Scientific 0.075 × 250mm, particle size 3 μm and pore size 100 Å). Prior to separation, the peptides were concentrated on a C18 precolumn (Thermo Scientific 0.1 × 50mm, particle size 5 μm and pore size 100 Å). Mobile phases were 100% water 0.1% formic acid (FA) (buffer A) and 100% Acetonitrile 0.1% FA (buffer B). Column gradient was developed in a gradient from 2% B to 40% B in 120 min. Column was equilibrated in 95% B for 10 min and 2% B for 10 min. During all process, precolumn was in line with column and flow maintained all along the gradient at 300 nl/min. Output of the separation column was directly coupled to nano-electrospray source. MS1 spectra was collected in the range of 350-1250 m/z for 250 ms. The 35 most intense precursors with charge states of 2 to 5 that exceeded 150 counts per second were selected for fragmentation, rolling collision energy was used for fragmentation and MS2 spectra were collected in the range of 230-1500 m/z for 100 ms. The precursor ions were dynamically excluded from reselection for 15 s. MS/MS data acquisition was performed using AnalystTF 1.7 (Sciex) and spectra files were processed through ProteinPilot v5.0 search engine (Sciex) using Paragon™ Algo-rithm (v.4.0.0.0) for database search. To avoid using the same spectral evidence in more than one protein, the identified proteins were grouped based on MS/MS spectra by the Progroup™ algorithm, regardless of the peptide sequence assigned. The protein within each group that could explain more spectral data with confidence was depicted as the primary protein of the group. FDR was performed using a non-lineal fitting method (Tang et al., 2008) and displayed results were those reporting a 1% Global FDR or better.

#### 2.5.3 SWATH-mass spectrometry proteomics: Quantitative analysis

Protein extracts (20 μg) from each sample were reduced by addition of DTT to a final concentration of 10mM and incubation at room temperature for 30 minutes. Subsequent alkylation by 30 mM iodoacetamide was performed for 30minutes in the dark. An additional reduction step was performed by 30mM DTT, allowing the reaction to stand at room temperature for 30 min. The mixture was diluted to 0.6M urea using MilliQ-water, and after trypsin addition (Promega) (enzyme:protein, 1:50 w/w), the sample was incubated at 37°C for 16h. Digestion was quenched by acidification with acetic acid. The digestion mixture was dried in a SpeedVac. Purification and concentration of peptides was performed using C18 Zip Tip Solid Phase Extraction (Millipore). The peptides recovered were reconstituted into a final concentration of 0.5μg/μL of 2% ACN, 0.5% FA, 97.5% MilliQ-water prior to mass spectrometric analysis. For SWATH-MS-based experiments the instrument (Sciex TripleTOF 5600+) was configured as described by Gillet et al (Gillet et al., 2012). Briefly, the mass spectrometer was operated in a looped product ion mode. In this mode, the instrument was specifically tuned to allow a quadrupole resolution of Da/mass selection. The stability of the mass selection was maintained by the operation of the Radio Frequency (RF) and Direct Current (DC) voltages on the isolation quadrupole in an independent manner. Using an isolation width of 16 Da (15 Da of optimal ion transmission efficiency and 1 Da for the window overlap), a set of 37 overlapping windows were constructed covering the mass range 450-1000 Da. In this way, 1 μL of each sample was loaded onto a trap column (Thermo Scientific 0.1 × 50mm, particle size 5 μm and pore size 100 Å) and desalted with 0.1% TFA at 3 μL/min during 10 min. The peptides were loaded onto an analytical column (Thermo Scientific 0.075 × 250mm, particle size 3 μm and pore size 100 Å) equilibrated in 2% acetonitrile 0.1% FA. Peptide elution was carried out with a linear gradient of 2 to 40% B in 120 min (mobile phases A:100% water 0.1% formic acid (FA) and B: 100% Acetonitrile 0.1% FA) at a flow rate of 300 nL/min. Eluted peptides were infused in the mass spectrometer. The Triple-TOF was operated in swath mode, in which a 0.050 s TOF MS scan from 350 to 1250 m/z was performed, followed by 0.080 s product ion scans from 230 to 1800 m/z on the 37 defined windows (3.05 s/cycle). Collision energy was set to optimum energy for a 2 + ion at the center of each SWATH block with a 15 eV collision energy spread. The mass spectrometer was always operated in high sensitivity mode. The resulting ProteinPilot group file from library generation was loaded into PeakView^®^ (v2.1, Sciex) and peaks from SWATH runs were extracted with a peptide confidence threshold of 99% confidence (Unused Score ≥ 1.3) and a FDR lower than 1%. For this, the MS/MS spectra of the assigned peptides was extracted by ProteinPilot, and only the proteins that fulfilled the following criteria were validated: (1) peptide mass tolerance lower than 10 ppm, (2) 99% of confidence level in peptide identification, and (3) complete b/y ions series found in the MS/MS spectrum. Only proteins quantified with at least two unique peptides were considered. The quantitative data obtained by PeakView® were analyzed using Perseus software 1.6.14 version (Tyanova et al., 2016) for statistical analysis and data visualization. MS data and search results files were deposited in the Proteome Xchange Consortium via the JPOST partner repository (https://repository.jpostdb.org) (Okuda et al., 2017) with the identifier PXD027645 for ProteomeXchange and JPST001274 for jPOST (for reviewers: https://repository.jpostdb.org/preview/6417285016102b4b16aaa0; Access key: 3355). Interactome and pathway analysis were performed using Metascape (Zhou et al., 2019) and machine learning based bioinformatic QIAGEN’s Ingenuity Pathway Analysis (IPA, QIAGEN Redwood City, www.qiagen.com/ingenuity)

### 2.6 Protein arrays

For the secretome analysis, a dot-blot protein array was used for cytokine profiling (Abcam). Briefly, membranes with 62 cytokine antibodies were blocked with the manufacturer’s blocking bu□er at room temperature (RT) for 30 min, and incubated o/n with 1 ml of undiluted cell-cultured media from OBC1 transfected cells (n = 3/condition). After washing, a biotinylated anti-cytokine antibody mixture was added to the membranes followed by incubation with HRP-conjugated streptavidin and then exposed to the manufacturer’s peroxidase substrate.

### 2.7 Western-blotting

Equal amounts of protein (10 μg) were resolved in 4-15% stain free SDS-PAGE gels (BioRad). Protein extracts derived from OBC1 cells were electrophoretically transferred onto nitrocellulose membranes using a Trans-blot Turbo transfer system (up to 25V, 7min) (BioRad). Membranes were probed with primary antibodies at 1:1000 dilution in 5% nonfat milk or BSA according to manufacturer instructions. After incubation with the appropriate horseradish peroxidase-conjugated secondary antibody (1:5000), the immunoreactivity was visualized by enhanced chemiluminiscence (Perkin Elmer) and detected by a Chemidoc MP Imaging System (Bio-Rad). Equal loading of the gels was assessed using Stain-free imaging technology. Thus, protein normalization was performed by measuring total protein directly on the gels used for western blotting. After densitometric analyses (Image Lab Software Version 5.2; Bio-Rad), optical density values were expressed as arbitrary units and normalized to total protein levels.

## 3. Results and discussion

Multiple studies have pointed out that SARS-CoV-2 infection affects the chemosensory processing in animals and humans, especially the olfaction. SARS-CoV-2 is able to reach the OE and the OB in mice and golden Syrian hamsters (de Melo et al., 2021;Ye et al., 2021). In rhesus monkeys, SARS-CoV-2 also invades the CNS primarily via the OB (Jiao et al., 2021). Although it has been proposed that the olfactory nerve is not a likely route to brain infection in COVID-19 patients (Butowt et al., 2021), several reports have shown the presence of SARS-CoV-2 (mRNA/protein levels or viral particles) in multiple brain areas including the OB derived from COVID-19 patients (Deigendesch et al., 2020;Matschke et al., 2020;Menter et al., 2020;Morbini et al., 2020;Lopez et al., 2021;Lou et al., 2021;Meinhardt et al., 2021;Serrano et al., 2021). In particular, the S protein has the capacity to cross the blood-brain barrier and reach the OB in mice (Rhea et al., 2021). In view of these preclinical and neuropathological data, an in-depth olfactory molecular characterization is necessary to unveil the missing links in the mechanisms of the sudden smell impairment induced by SARS-CoV-2 infection. To address this gap in knowledge, we have evaluated in-depth the OB cell response upon the intracellular expression of the SARS-CoV-2 structural proteins. The S glycoprotein is the main determinant of viral entry in target cells (Rhea et al., 2021). The M glycoprotein is key for viral particle assembly (Escors et al., 2001;de Haan and Rottier, 2005), whereas the E protein is a multifunctional protein, acting as a viroporin in its oligomeric form, participating on viral assembly and virion release (Pervushin et al., 2009;Schoeman and Fielding, 2019). The N protein packages the positive strand viral genome RNA into a helical ribonucleocapsid (RNP), playing a fundamental role during virion assembly through its interactions with the viral genome and membrane protein M (Escors et al., 2001;Satarker and Nampoothiri, 2020). In general, SARS-CoV-2 structural protein factors are essential in the viral genome replication, receptor attachment and virion formation, promoting the viral spreading and pathogenesis.

### 3.1 Molecular derangements induced by SARS-CoV-2 structural proteins in OB cells

To examine the molecular alterations due to the presence of SARS-CoV-2 structural proteins in an olfactory cell system, GFP-tagged SARS-CoV-2 structural proteins S, M, N and E were expressed in OBC1 cells (**Figure 1A**). After GFP-positive cell sorting, the olfactory metabolic imbalance induced by each SARS-CoV-2 protein was monitored across all experimental conditions by RNA-seq and shotgun proteomics. At transcriptome level, 1506, 1440, 359 and 1237 differential expressed genes were found between control-GFP and GFP-E, GFP-M, GFP-N and GFP-S SARS-CoV-2 proteins respectively (p-value adjusted <0.01; fold-change: 50%) (**Supplementary Table 1**). In contrast, around 15-23% of the quantified proteome was modulated by GFP-tagged SARS-CoV-2 structural proteins. Specifically, 435, 387, 594 and 584 proteins differential expressed proteins (DEPs) were found between Control-GFP and GFP-S, GFP-N, GFP-M and GFP-E respectively (p-value <0.05; fold-change 30%) (**Supplementary Table 2**). GFP-tagged SARS-CoV-2 proteins were detected by Western-blotting (**Supplementary Figure 1A**) and mass-spectrometry (**Supplementary Table 2**) except the GFP-E protein, due to its small size and the high content of aliphatic hydrophic aminoacids in its sequence (36% of Leu and Val) (Duart et al., 2020). Multiple commonalities and differences were observed when transcriptomic and proteomic datasets derived from the expression of GFP-tagged SARS-CoV-2 structural proteins were compared (**Figure 1B-C; Supplementary figures 1B, 2A**). Focusing on the most affected proteins, we identified a downregulation in Ak4 (Adenylate kinase 4), a protein that controls ATP levels through the regulatory activity of AMPK which plays a protective role against oxidative stress (Liu et al., 2009;Lanning et al., 2014). A detailed functional clustering revealed that SARS-CoV-2 structural proteins independently impact on multiple pathways at transcriptomic and proteomic levels (**Supplementary figures 1C, 2B; Supplementary Tables 3-4**). Comparing the functionality derived from both omic approaches, all SARS-CoV-2 structural proteins commonly interferes with multiple biofunctions such as extracellular matrix organization, deregulation of lipid metabolism, hemostasis, non-integrin membrane interactions and signaling by receptor tyrosine kinases (**Figure 1D**). However, SARS-CoV-2 structural proteins commonly induced specific alterations with different impact on transcriptomic homeostasis and proteostasis. Considering the most statistically significant enriched pathways, SARS-CoV-2 proteins interfere with transcriptional activities associated to chondroitin/dermatan sulphate metabolism, platelet homeostasis, PPAR-dependent gene expression, opioid signaling, glycosylation and cytokine, IGF and PDGF signaling (**Figure 1D**). At proteostatic level, SARS-CoV-2 structural proteins modulate aminoacid metabolism, biological oxidations, interleukin-12 signaling and processing of pre-mRNA and rRNA, between others (**Figure 1D**).

**Figure 1.**
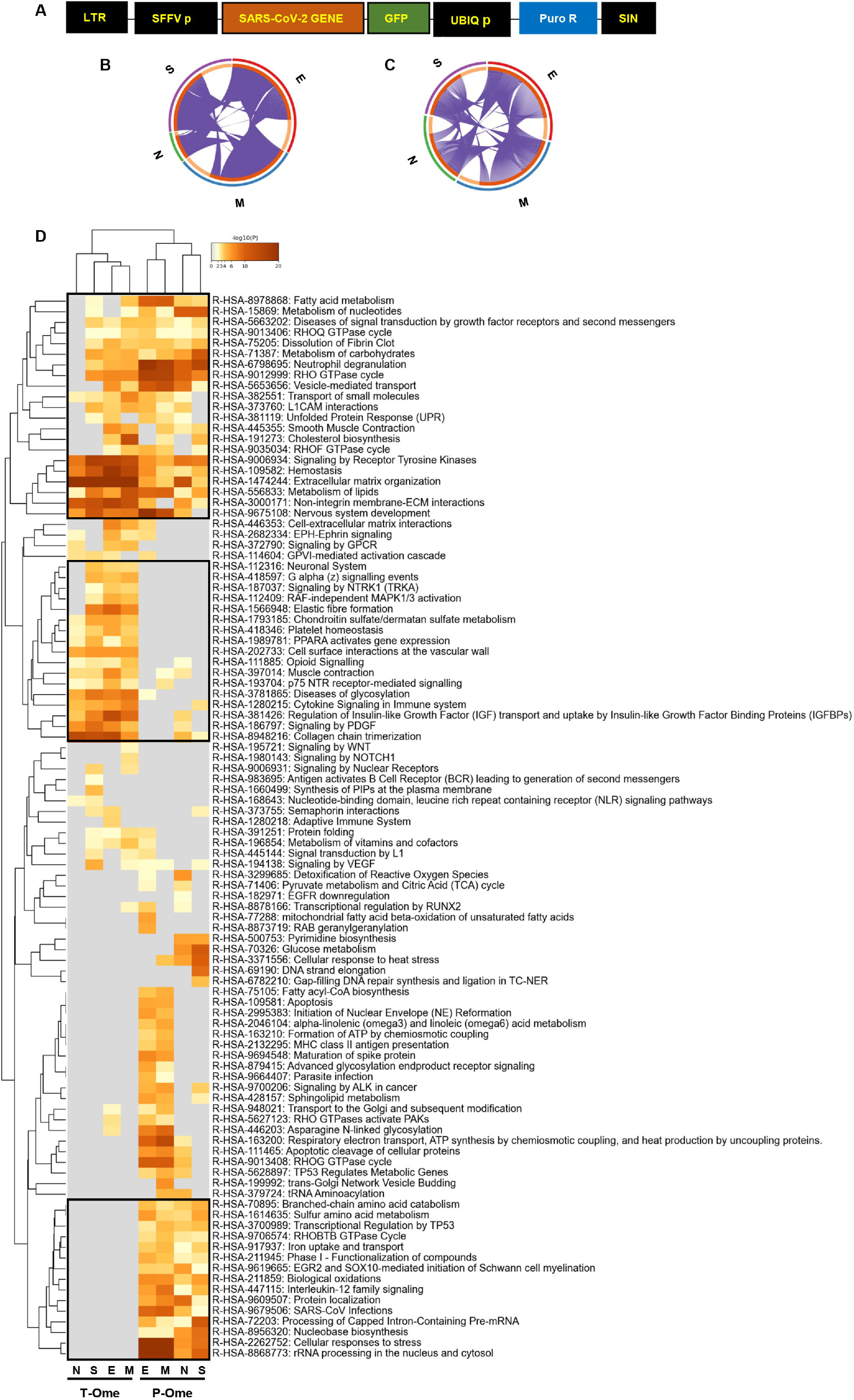
Molecular dyshomeostasis induced by SARS-CoV-2 structural proteins in OBC1 cells. A) Lentivector expression vectors used in this study. LTR, long terminal repeats; SFFV, spleen focus-forming virus promoter; GFP-tagged SARS-CoV-2 structural proteins (M, N, E, S); UBIQp, ubiquitin promoter; PuroR, gene conferring puromycin resistance. B and C) Circus plot of dysregulated transcripts (B) and proteins (C) targeted by each SARS-CoV-2 structural proteins. Purple lines indicate the common deregulated transcriptome/proteome across OBC1 cells that express each GFP-tagged SARS-CoV-2 structural protein. D) Functional clustering representing the significantly altered pathways by SARS-CoV-2 structural proteins at the transcriptome (T-Ome) and proteome (P-Ome) layers. After the identification of all statistically enriched Reactome terms, cumulative hypergeometric p values and enrichment factors were calculated and used for filtering. Remaining significant terms were then hierarchically clustered into a tree based on Kappa-statistical similarities among their gene memberships. 0.3 kappa score was applied as the threshold to cast the tree into term clusters. The term with the best p value within each cluster was selected as its representative term and displayed in a dendrogram. The heat map cells are coloured by their p values; white cells indicate the lack of enrichment for that term in the corresponding protein/gene list. Squares (black frame) highlight significantly enriched terms detected at transcript, protein levels or both.

To decipher specificities and singularities associated to the expression of each SARS-CoV-2 structural protein at olfactory level, functional analyses were performed excluding the common deregulated proteome and taking into account only the mapping of unique DEPs associated to each SARS-CoV-2 protein (**Supplementary Table 5**). Interestingly, densely connected complexes with regulatory roles in GTP hydrolysis, calcium uptake into mitochondria (processing SMDT1), nuclear pore complex formation, vesicle-mediated transport, antigen presentation and peroxisomal protein import were commonly altered by structural SARS-CoV-2 proteins (**Figure 2,** upper). However, protein components of these complexes were differentially targeted by each structural viral protein (**Figure 2,** lower). Inferring more biologically interpretable results, GFP-M significantly modulated protein mediators located in organelle membranes that are mainly involved in trans-Golgi network vesicle budding, ferroptosis and cellular response to chemical stress (**Figure 3A and B**). GFP-S independently impacted on nuclear proteins regulating cholesterol synthesis, DNA elongation and mRNA splicing between others (**Figure 3A and B**). GFP-E-dysregulated proteins presented a widespread localization, affecting a plethora of non-related functions such as mRNA processing, RAB geranylgeranylation and integrin interactions, whereas GFP-N specifically altered lipid, glutathione and glyoxilate metabolism (**Figure 3A and B**). In contrast to recent results with full viral particles, these data generated at the organelle-complex-pathway axis demonstrate the enriched multifactorial functions associated to each SARS-CoV-2 protein. Interestingly, 14-18% of the perturbations generated by SARS-CoV-2 M, N and E proteins (at the level of protein-coding gene) in OBC1 cells were common with data derived from pulmonary cells expressing these structural viral proteins (Stukalov et al., 2021) (**Figure 3C**). However, only 3% of the proteome modulated by the S protein was shared across studies, indicating that a great proportion of proteostatic events intercepted by structural SARS-CoV-2 proteins are cell-dependent. In addition, part of the altered molecular routes identified in our in-vitro system have been recently detected in the OB from COVID-19 subjects (Piras et al., 2021) (**Supplementary Figure 3**), pointing out specific pathways directly targeted by SARS-CoV-2 structural proteins beyond the CNS affectation and/or the systemic immune response to SARS-CoV-2 infection.

**Figure 2.**
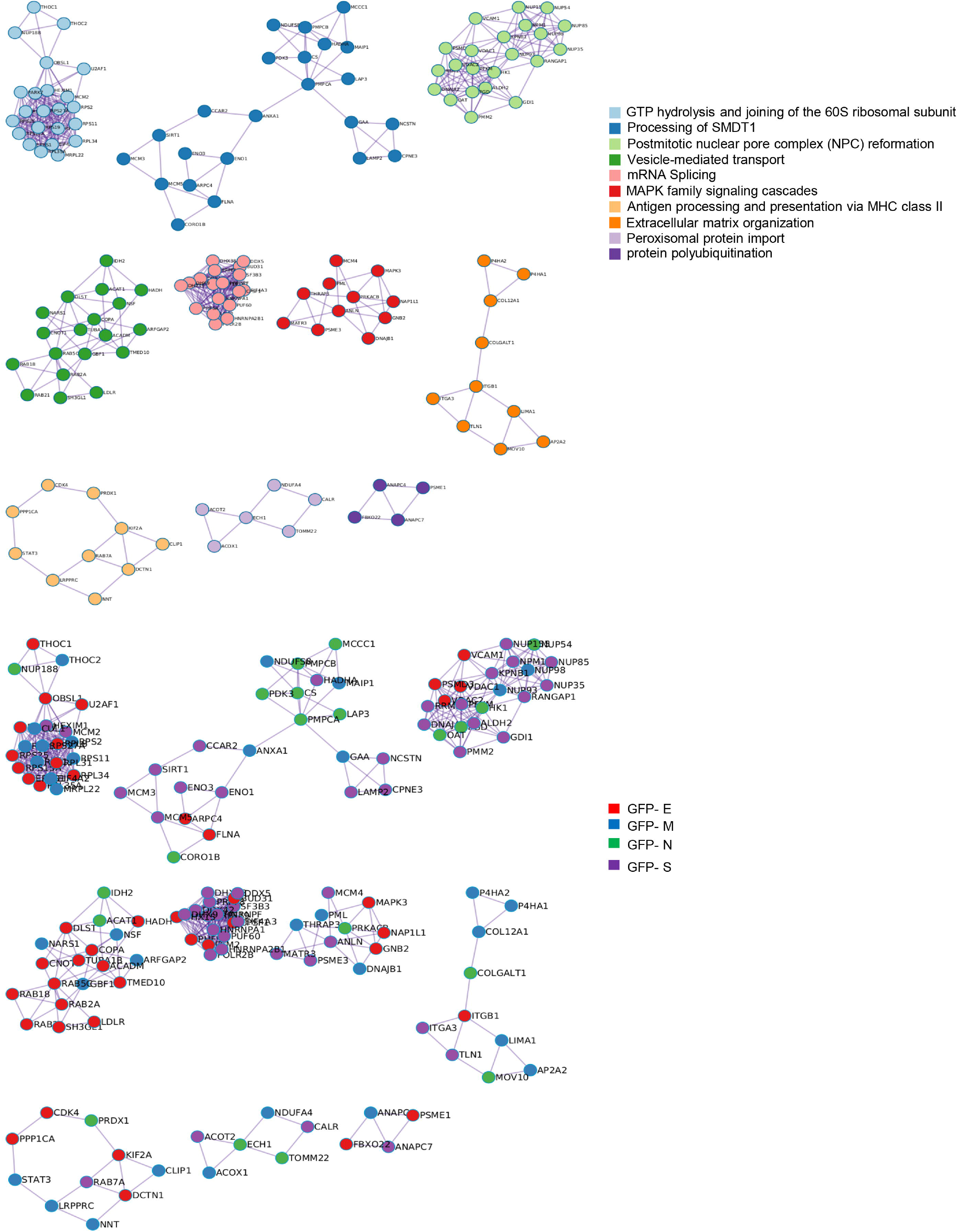
Intracellular protein complexes affected by SARS-CoV-2 structural protein expression. MCODE algorithm (Bader and Hogue, 2003) was applied to automatically extract protein complexes embedded in proteomics datasets. The three most significantly enriched ontology terms were combined to annotate putative biological roles for each MCODE complex (Upper). Each GFP-tagged SARS-CoV-2 protein differentially modulated each MCODE complex in OBC1 cells (lower).

**Figure 3.**
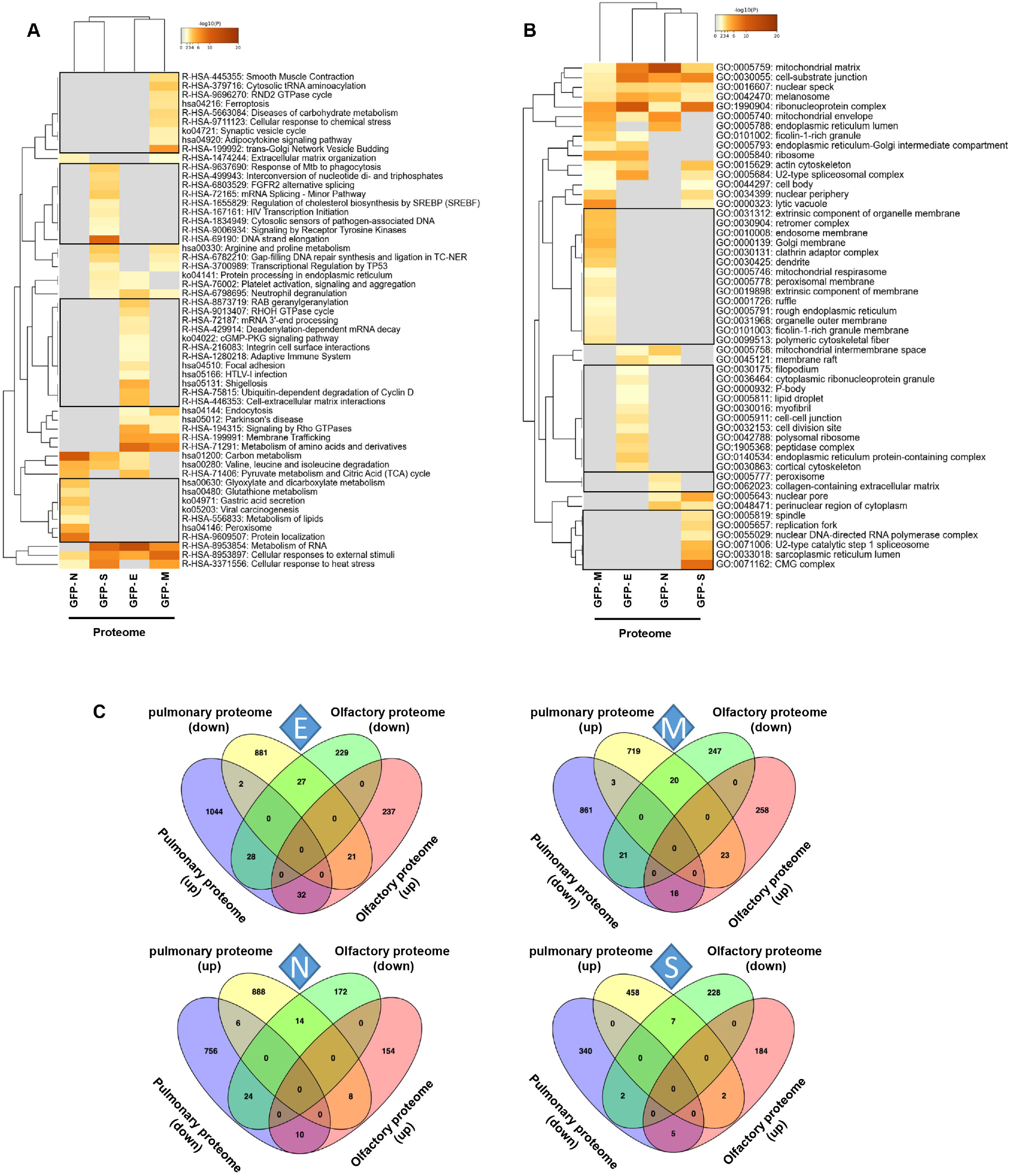
Functional clustering based on the specific OBC1 proteome disrupted by each SARS-CoV-2 structural protein. A) Reactome-based pathway enriched clusters across diferential proteomes induced by the expression of GFP-SARS-CoV-2 structural proteins. B) GO ontology clustering focus on subcellular compartments. Squares (black frame) represent significantly enriched terms derived from the independent expression of each SARS-CoV-2 structural protein. C) Commonalities and differences between proteostatic changes induced by the expression of SARS-CoV-2 structural proteins in olfactory OBC1 and pulmonary A549 cells (Stukalov et al., 2021).

### 3.2 SARS-CoV-2 structural proteins differentially modulate the activation profile of survival routes and mitochondrial homeostasis in OB cells

To enhance the analytical outcome of proteostatic alterations, proteome-scale interaction networks were constructed merging the differential proteomes induced by SARS-CoV-2 structural proteins. As shown in **Figure 4**, both p38 MAPK and Akt appeared as principal nodes in protein interactome maps. Although much effort has been spent on characterizing the pleiotropic effects of SARS-CoV-2 infection in multiple biological contexts, there is no detailed information about the impact of each structural SARS-CoV-2 protein on the activation dynamics of the cell survival. Even though changes in p38 MAPK and Akt expression were not detected in our proteomic analysis, the alteration of some targets might be indicative of a dysfunctional state when SARS-CoV-2 structural proteins are continuously expressed. Subsequent experiments were performed to analyze the activation state of p38 MAPK and Akt signaling pathways across GFP-SARS-CoV-2 structural protein expressing-OBC1 cells (**Figure 5**). As shown in **Figure 5A**, the presence of SARS-CoV-2 structural proteins induced a p38 MAPK and Akt inactivation. Specifically, the independent expression of SARS-CoV-2 S and N proteins inactivated both kinases. In contrast, the M protein specifically inactivated Akt and the E protein interfered with p38 MAPK. To complement our signaling mapping, other stress-responsive kinases relevant to OB homeostasis (Lachen-Montes et al., 2019) were checked. N and S proteins also induced the inactivation of GSK3 and the SEK1 stress-activated protein kinase (SAPK) axis (**Figure 5B**). However, S protein expression induced the inactivation of PKA whereas the expression of the N protein enhanced the active phosphorylation of the PKA catalytic subunit (**Figure 5B**). No appreciable changes were observed in the activation profile of other survival kinases such as PDK1, PKC, CaMKII and MEK1/2 (**Supplementary Figure 4**). These data demonstrate that SARS-CoV-2 structural proteins differentially impact on the survival potential of OB cells. Many viral infections trigger the activation of the p38 MAPK and Akt signaling pathways for an efficient replication, including SARS-CoV-2 (Kindrachuk et al., 2015;Appelberg et al., 2020;Cheng et al., 2020). p38 MAPK and Akt inactivation induced by the expression of SARS-CoV-2 structural proteins could be part of an antiviral mechanism triggered in olfactory cells. In fact, pharmacological inhibition of p38 MAPK has antiviral efficacy (Bouhaddou et al., 2020;Grimes and Grimes, 2020) and Akt inhibitors have been proposed as candidate drugs (Fagone et al., 2020). Therefore, our results are in agreement with evidence pointing to both kinases as potential therapeutic targets for COVID-19. GSK3 is essential for N phosphorylation and SARS-CoV-2 replication (Liu et al., 2021). Similarly to SARS-CoV-1, GSK3 activity may be also critical for the initiation of oxidative stress, and inflammation during SARS-CoV-2 infection (Rana et al., 2021), raising the possibility that targeting GSK-3 could open new therapeutic opportunities in COVID-19 (Rudd, 2020). SEK1 (MKK4) is an essential component of the stress-activated protein kinase/c-Jun N-terminal kinase (SAP/JNK) signaling pathway that is phosphorylated by the avian coronavirus infectious bronchitis virus (IBV) (Fung and Liu, 2017). However, the mechanistic connection with SARS-CoV-2 is not known. We observed an enhanced PKA activity when SARS-CoV-2 N protein is overexpressed in OBC1 cells. Interestingly, the co-expression of PKA and N protein generates a poliphosphorylated N dimers able to sequestrate cellular 14-3-3 isoforms (Tugaeva et al., 2021).

**Figure 4.**
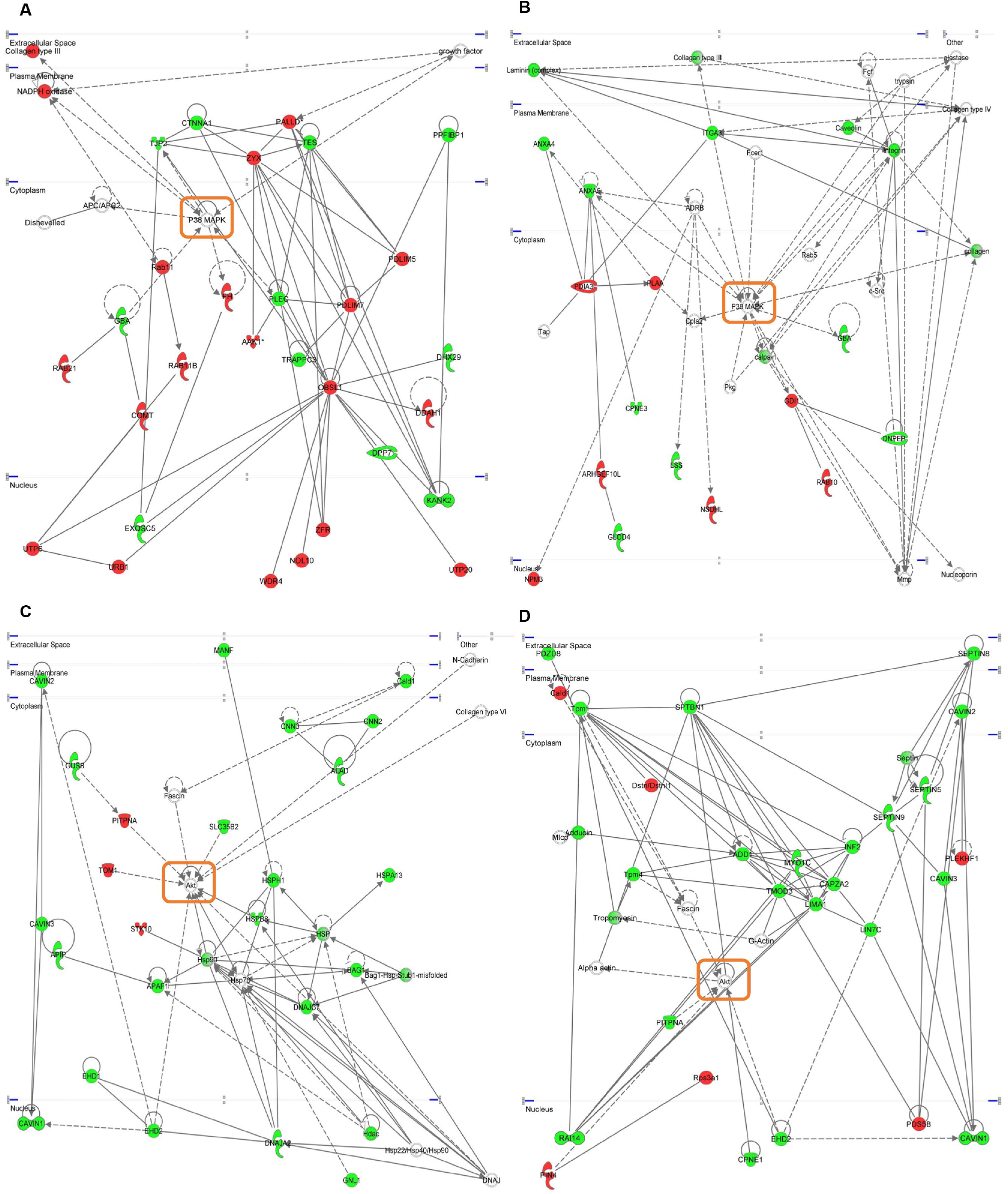
Representative functional protein interactome maps for differentially expressed proteins in GFP-SARS-CoV-2 structural protein-expressing OBC1 cells. Visual representation of the functional relationships between altered proteome generated by the expression of SARS-CoV-2 structural E protein (A), S protein (B,C) and M protein (D) in OBC1 cells. Dysregulated proteins are highlighted in red (up-regulated) and green (down-regulated). Orange circles point the principal nodes of the interactome maps. Continuous and discontinuous lines represent direct and indirect interactions respectively.

**Figure 5.**
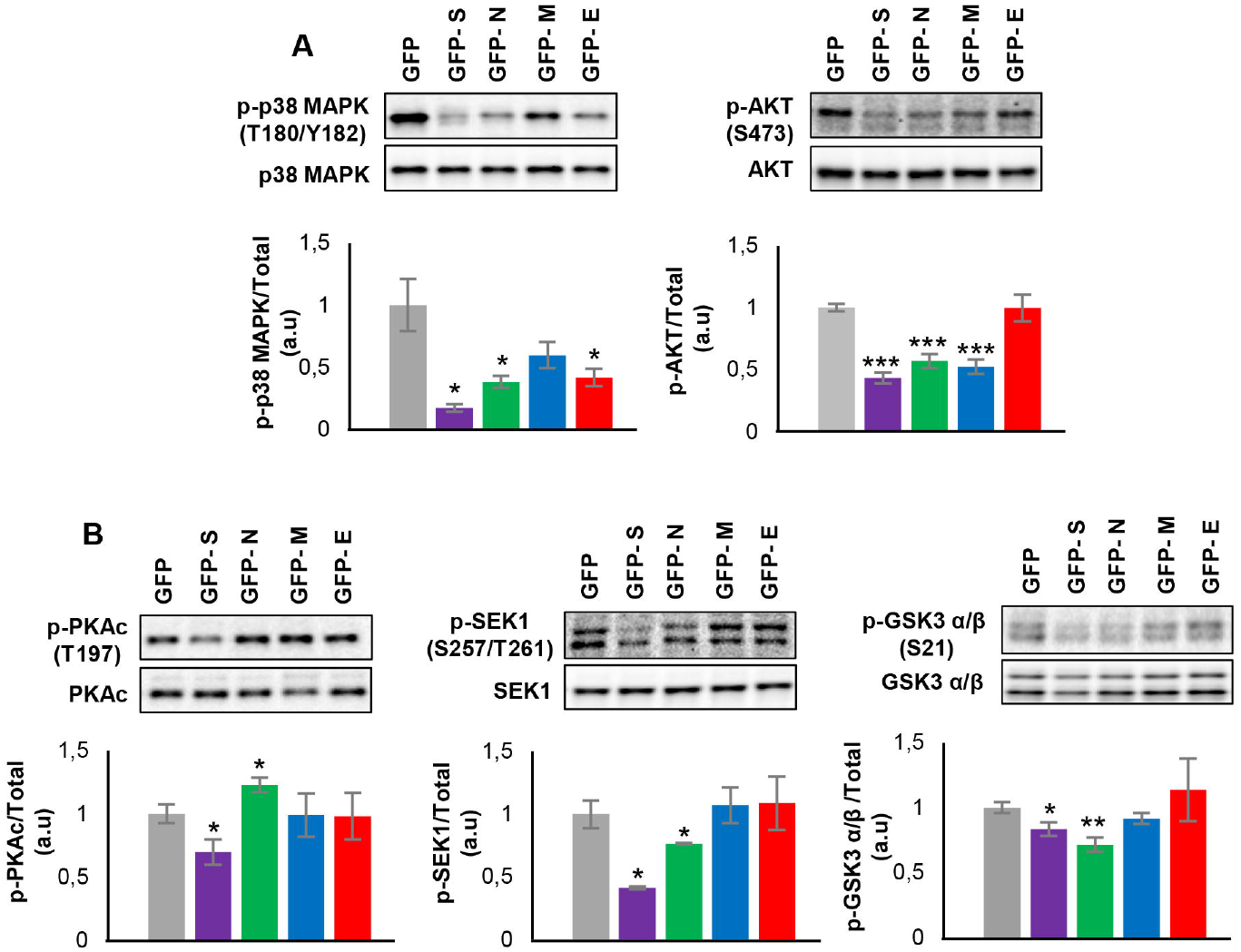
Levels and residue-specific phosphorylation of A) p38 MAPK, Akt and B) PKA, SEK1 and GSK3 in GFP-SARS-CoV-2 structural protein expressing-OBC1 cells. Equal loading of the gels was assessed by stain-free digitalization. Panels show histograms of band densities. Data are presented as mean ± SEM from four independent biological replicates. *P < 0.05 vs. GFP; **P < 0.01 vs. GFP; *** P < 0.001 vs. GFP (a.u: arbitrary units).

It has been recently demonstrated the existence of physical and functional interactions between SARS-CoV-2 and host mitochondria, serving this organelle as a subcellular platform for anti-SARS-CoV-2 immunity (Flynn et al., 2021). Although mitochondrial proteostasis was modified by all SARS-CoV-2 structural proteins (**Figure 3B**), our network analysis uncovered a generalized down-regulation of mitochondrial complex I subunits specifically induced by the expression of SARS-CoV-2 E and M proteins (**Figure 6A, B**). Mitochondrial prohibitin complex (constituted by Phb1 and Phb2 subunits) modulates mitochondrial dynamics and respiratory complex assembly, triggering beneficial effects by reducing free radical production (Artal-Sanz and Tavernarakis, 2009;Zhou et al., 2012). According to Biogrid database (https://thebiogrid.org/), Phb complex interacts with SARS-CoV-2 proteome at multiple levels. Specifically, Phb1 interacts with ORF9B/C, NSP2, NSP3, NSP6 and NSP8 whereas NSP2, NSP3, NSP6, ORF6, ORF9B, E and M proteins are Phb2 interactors. As shown in **Figure 6C**, a depletion in Phb1 levels was observed in E protein expressing OBC1 cells. In contrast, a significant increment in Phb2 levels was evidenced when S and N proteins were produced. In general, Phb1 repression triggers a concomitant reduction of its partner Phb2 and viceversa (Merkwirth et al., 2012), indicating that SARS-CoV-2 structural proteins has the capacity to impact on mitochondrial homeostasis differentially interfering with the functional interdependency of Phb subunits.

**Figure 6.**
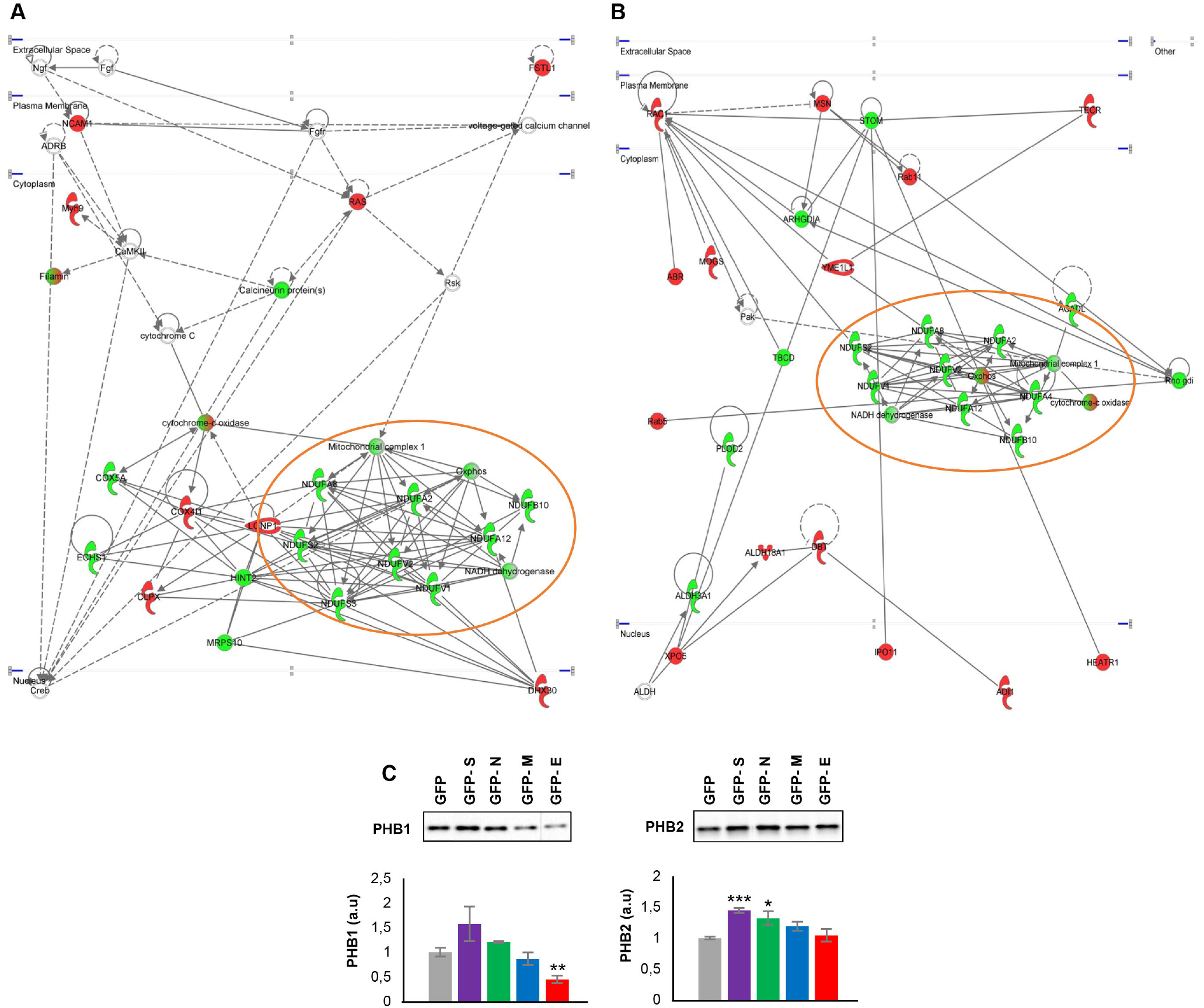
Functional protein interactome maps for differentially expressed proteins in GFP-SARS-CoV-2 E and M protein-expressing OBC1 cells, revealing a mitochondrial impairment. (A, B). OB expression of Phb complex by Western blot (C). Equal loading of the gels was assessed by stain-free digitalization. Panels show histograms of band densities. Data are presented as mean ± SEM from four independent biological replicates. *P < 0.05 vs. GFP; **P < 0.01 vs. GFP; *** P < 0.001 vs. GFP (a.u: arbitrary units).

### 3.3 SARS-CoV-2 structural proteins induce a dissimilar immunological effectome in OB cells

The vast amount of information about the relationship between immune system and COVID-19 points out that multiple aberrations of innate and acquired immunity are induced by SARS-CoV-2 infection. Specifically, astrogliosis, microgliosis and infiltration by cytotoxic T lymphocytes in the OB have been characterized in COVID-19 subjects (Matschke et al., 2020). This olfactory neuroinflammation is different from the observed in other brain areas (Schwabenland et al., 2021). In addition, it has been proposed that olfactory ensheating cells may produce glia transit tubules through which cytokines and chemokines might migrate across the OE-OB axis (Xydakis et al., 2021). However, little is known about the immunological mechanisms triggered by each SARS-CoV-2 structural protein at olfactory level. In agreement with previous reports (Stukalov et al., 2021), specific gene/protein subsets modulated by each SARS-CoV-2 structural protein profile a pro-inflammatory signature (**Supplemental Table 6 and Supplementary Figure 5**), inducing specific changes in the OBC1 immunological effectome. As shown in **Figure 7**, S protein expression impacted on regulators of IL-6, IL-10 and IL-12 and in the differentiation of mature B cells. Alterations in protein-coding genes relevant in inflammasomes, lymphocyte migration, response to IL-7 and B cell activation were induced by the ex-pression of GFP-N protein expression in OBC1 cells (**Figure 7**). On the other hand, the M protein had the potential to interfere with lymphocyte activation, T cell differentiation, macrophage differentiation as well as in the IL-10, IL-27, and IL-35 signaling, whereas SARS-CoV-2 E protein specifically impacts on adaptive immune response, IL-17 pro-duction, B cell receptor signaling and natural killer cell differentiation between other processes (**Figure 7**).

**Figure 7.**
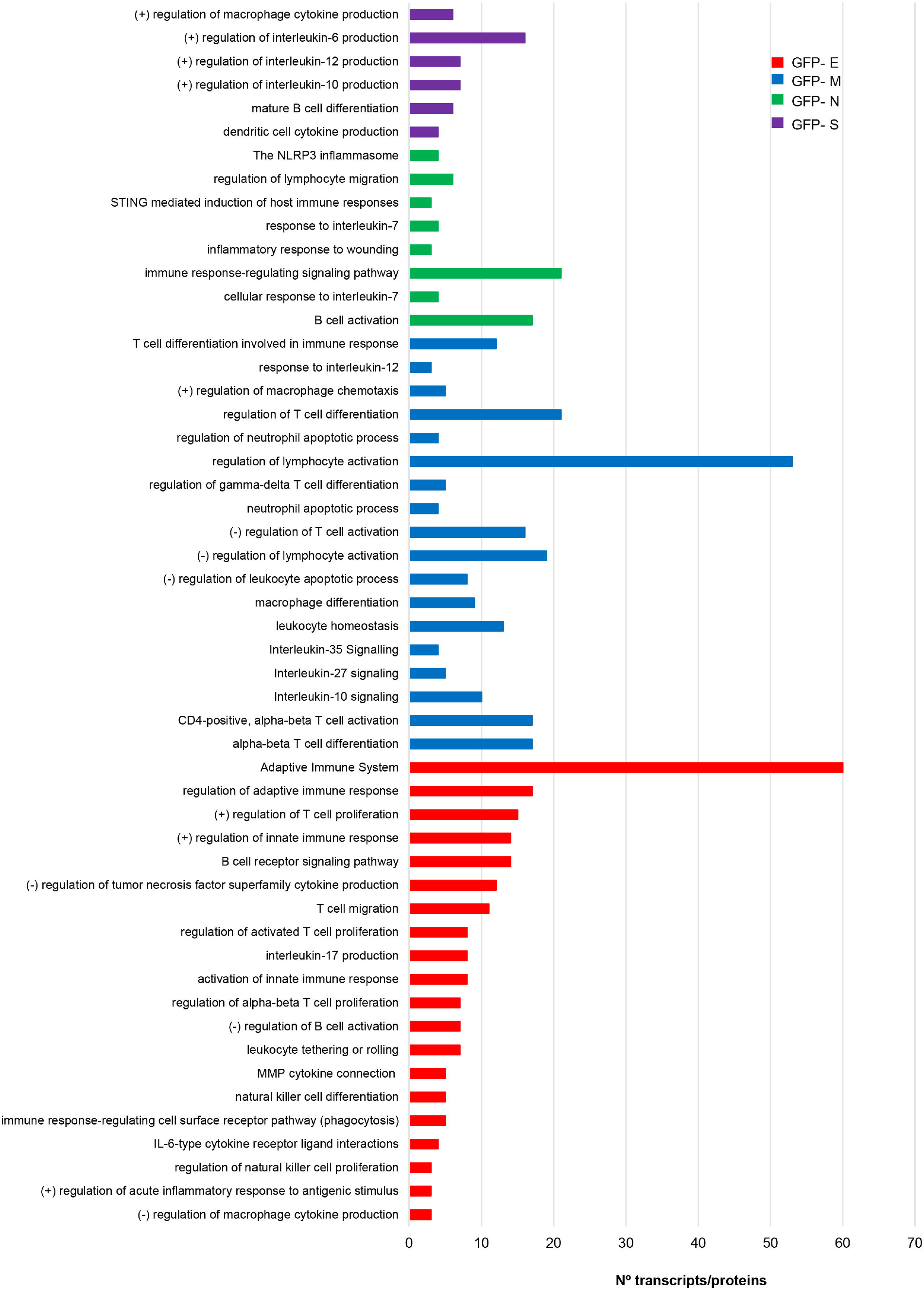
The expression of SARS-CoV-2 structural proteins differentially modulate the immunological capacity of OBC1 cells. Processes related to immune system were directly extracted from the functional analysis of proteotranscriptomic datasets (see **supplementary Table 6** for more details).

We wanted to partially map the consequences over the intracellular immunological effectome dependent on SARS-CoV-2 structural proteins. Cytokines and growth factors secreted by each stable OB cell line constitutively expressing a GFP-tagged SARS-CoV-2 structural protein may provide novel insights into the inflammatory imbalance produced by SARS-CoV-2. As shown in **Figure 8**, GFP-tagged M and E protein expression in OBC1 cells induced more extracellular changes. Part of the 62 secreted cell-cell signaling molecules analyzed were commonly altered between the structural proteins (**Figure 8**). Specifically, the increment of RANTES (CCL5) and MIP3a (CCL20) levels in the OBC1 secretome was independently induced by all structural proteins. Both chemokines are elevated in sera from COVID-19 patients (Hue et al., 2020). IFN gamma and MCP5 (CCL12) extracellular levels were also increased in SARS-CoV-2 S, M and E protein-expressing OBC1 cells (**Figure 8**). Although most of the differential soluble mediators have been previously related to the immune cell recruitment associated to COVID-19 cytokine storm (Ricci et al., 2021), these data contribute to the better understanding of the multifunctional immunomodulatory properties of SARS-CoV-2 M, N, S and E proteins beyond their intrinsic role in virion formation.

**Figure 8.**
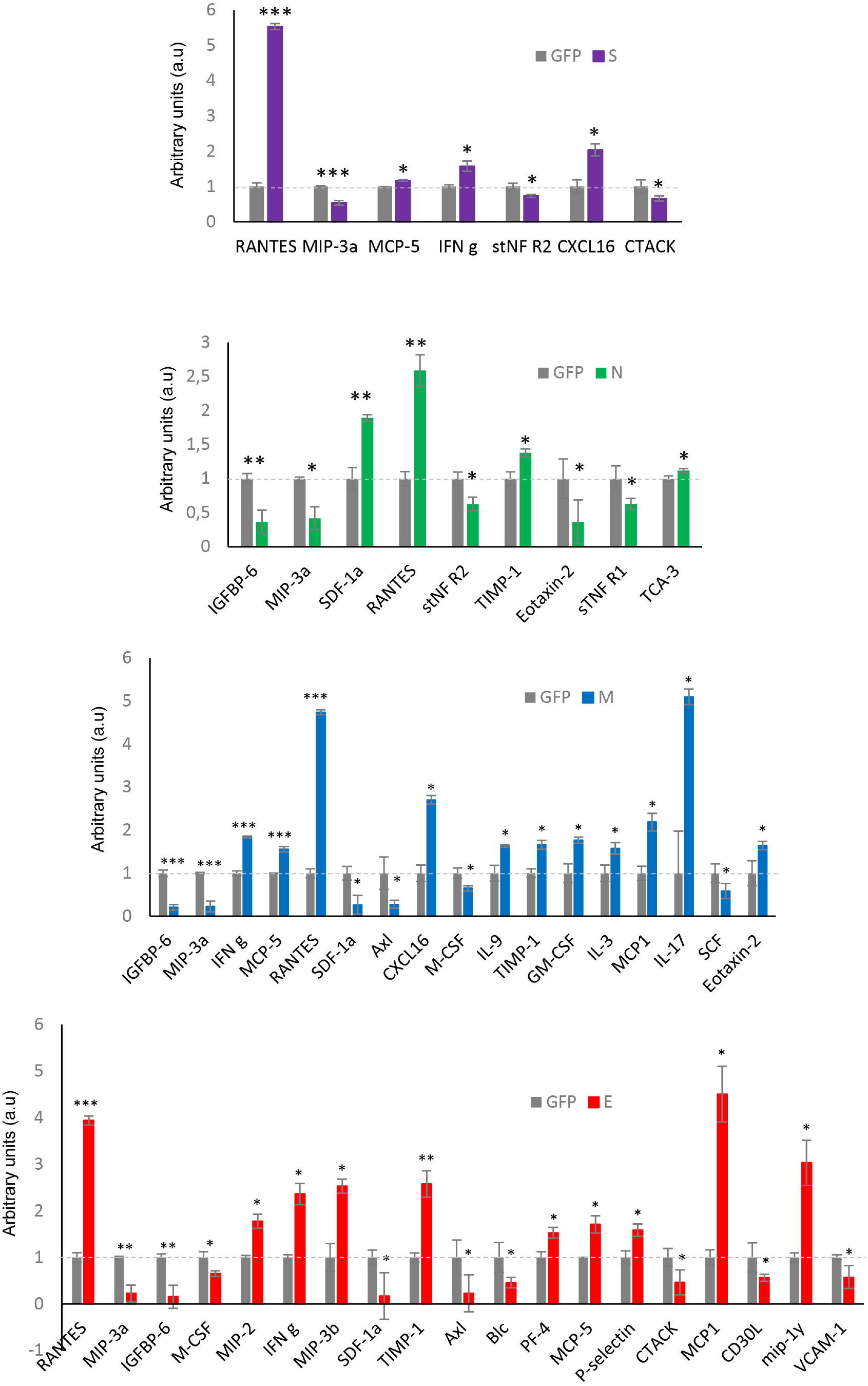
Extracellular cytokine profiling of GFP-SARS-CoV-2 structural protein-expressing OBC1 cells. The analysis of 62 cytokines/growth factors was performed in the cell media derived from all biological conditions using a dot-blot protein array method. Four independent experiments were performed. Data are presented as mean ± SEM. *p < 0.05, **p < 0.01, and ***p < 0.001 vs GFP.

### 3.4 Proteotranscriptomic data integration by machine learning unravels TGF-beta signaling route as a confluent activation node by SARS-CoV-2 structural proteins

It has been previously demonstrated that sophisticated machine learning (ML) approaches are useful workflows to integrate multi-omics data with the aim to discover complex interconnections between different type of entities (Ebrahim et al., 2016;Mann et al., 2021). To uncover regulatory complex composed by molecular effectors potentially modulated by SARS-CoV-2 structural proteins at olfactory level, transcriptomic and proteomic datasets were integrated and mined by a machine-learning approach in which multiple entities (genes, proteins, upstream regulators, pathways, diseases) were interconnected in the form of knowledge graphs. Considering the IPA Z□score as a statistical measure of the match between expected relationship direction and observed gene/protein expression, functional graphs were constructed. As shown in **Figure 9**, common hubs were predicted for each SARS-CoV-2 structural protein such as TGFB1/B3, SMAD and TEAD proteins, EGF, SPP1 and EDN1. It has been previously observed that SARS-CoV-2 N protein interacts with SMAD proteins enhancing TGF-beta signaling (Wang et al., 2022) whereas SARS-CoV-2 S protein triggers a transcriptional response associated to TGF-beta signaling (Biering et al., 2021). Interestingly, a therapeutic strategy focused on the inactivation of TGF-beta using integrin inhibitors has been proposed to mitigate COVID-19 severity (Huntington et al., 2022). EDN1 (Endothelin-1) has been considered a neuroprotective factor that participates in the olfactory response modulation through the uncoupling of gap junctions (Le Bourhis et al., 2014;Bryche et al., 2019). Interestingly, elevated EDN1 levels together with a sustained inflammation has been observed 3 months after recovery from acute COVID-19 symptoms (Willems et al., 2021). SPP1 (osteopontin) participates in the OB synaptic plasticity (Powell et al., 2019) and acts as a molecular brake regulating neuroinflammatory response to chronic viral infections (Mahmud et al., 2020). Involved in the enhancing production of interferon-gamma and interleukin-12, SPP1 is also overproduced in COVID-19 patients (Gibellini et al., 2020).

**Figure 9.**
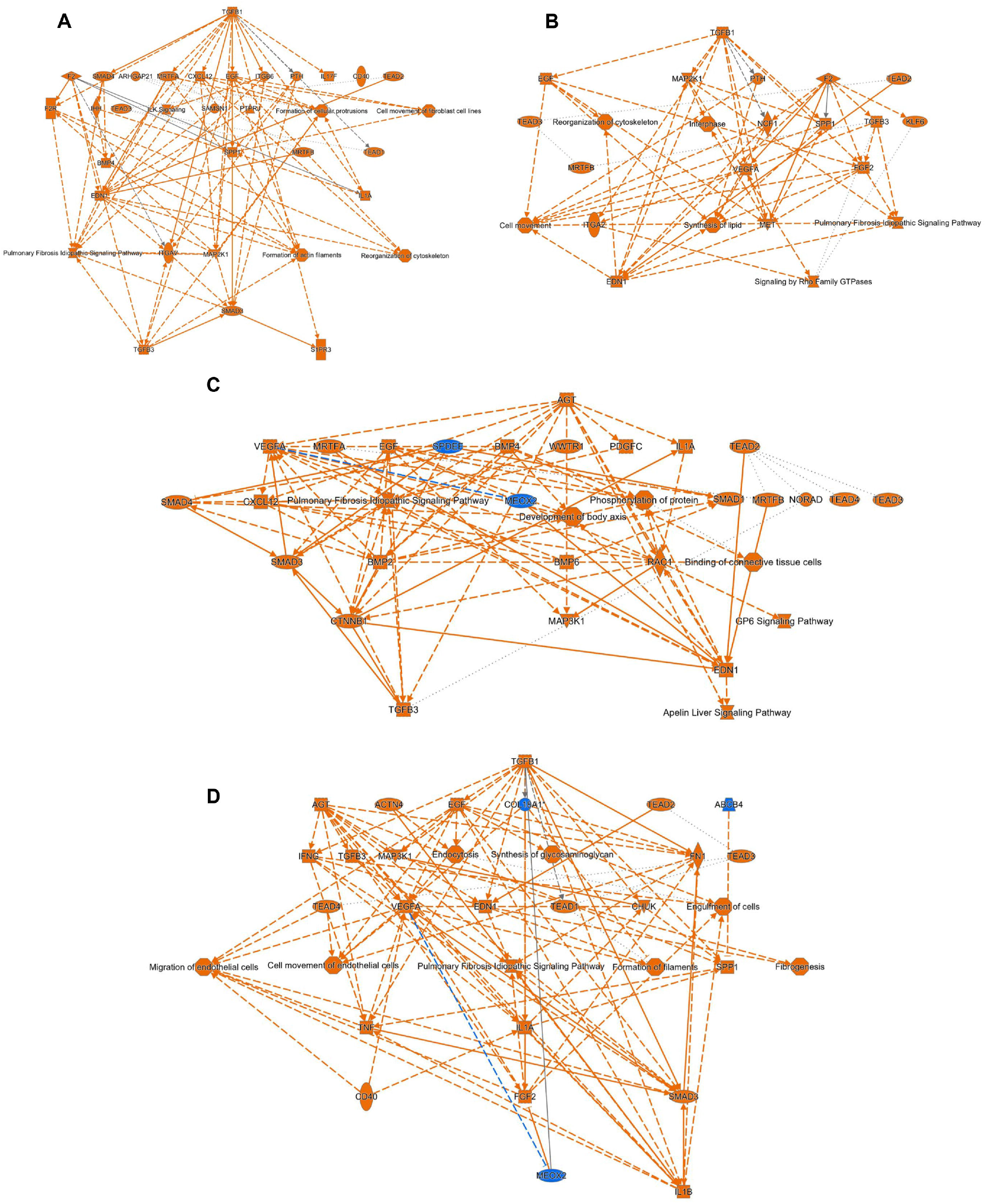
Machine learning-based analysis proposes a potential activation of TGFbeta/SMAD axis commonly induced by all SARS-CoV-2 structural proteins. The functional networks use a precomputed table containing inferred relationships between molecules, functions, diseases, and pathways obtained and scored by a machine learning algorithm operating entirely on prior knowledge. The heuristic graph algorithm present in IPA software was optimized to create a manageable network that brings together the most significantly activated (in orange; positive Z-score) or inhibited (in blue; negative z-score) upstream regulators, diseases, functions, and pathways from the differential proteotranscriptomic alterations induced by each SARS-CoV-2 structural proteins: A) SARS-CoV-2 E protein; B) SARS-CoV-2 M protein; C) SARS-CoV-2 N protein; D) SARS-CoV-2 S protein.

Although our study has uncovered the existence of commonalities and differences in the SARS-CoV-2 structural protein functionality, potential limitations exist that warrant discussion. It is important to note that SARS-CoV-2 structural and non-structural protein interactions are absent in our study (Boson et al., 2021;Xu et al., 2021). We are aware that using our multi-omic approach, the molecular dimension generated by posttranslational modifications (i.e phosphorylation) not only in host proteins but also in the SARS-CoV-2 structural proteins (Stukalov et al., 2021) has not been considered in this study, hampering the characterization of potential substrates modulated by the virus-hijacked kinase activation profiles. Due to our experimental design, the overexpression of exogenous proteins may generate drawbacks concerning protein misfolding, localization and regulation as well as intrinsic limitations associated to GFP expression systems (Jensen, 2012). Based on the olfactory cellular system used, additional experiments are needed to verify the specific role of the SARS-CoV-2 structural proteome in different human olfactory cellular contexts (Lachen-Montes et al., 2020;Hatton et al., 2021). As shown in previous reports performed at cellular and tissular levels (Nie et al., 2021;Stukalov et al., 2021), the application of proteomics in different olfactory cell layers as well as in olfactory areas directly derived from COVID-19 individuals, would increase our understanding of not only the early smell impairment associated to SARS-CoV-2 infection but also the olfactory recovery potential in COVID-19 patients.

## Supplementary Materials

**Supplementary Figure 1:** A) Expression of GFP-SARS-CoV-2 structural proteins. B) commonalities and differences across differential expressed proteins induced by SARS-CoV-2 proteome expression. C) Functional analysis of proteomics datasets.

**Supplementary Figure 2:** A) Commonalities and differences across differential expressed transcripts induced by SARS-CoV-2 proteome expression. B) Functional analysis of transcriptomics datasets.

**Supplementary Figure 3:** Comparative analysis between deregulated pathways observed in OBC1 cells expressing SARS-CoV-2 structural proteins and altered pathways observed at the level of the OB from COVID-19 subjects.

**Supplementary Figure 4:** Steady-state levels and activation state of PDK1, PKC, MEK1/2 and CaMKII in SARS-CoV-2 structural protein-expressing OBC1 cells.

**Supplementary Figure 5:** Differential immunological effectome (genes/proteins) induced by the expression of SARS-CoV-2 structural proteins in OBC1 cells.

**Supplementary Table 1:** OBC1 Differential transcriptome induced by each GFP-SARS-CoV-2 structural protein.

**Supplementary Table 2:** OBC1 Differential proteome induced by each GFP-SARS-CoV-2 structural protein.

**Supplementary Table 3:** Gene-Ontology (GO) functional analysis derived from transcriptomics.

**Supplementary Table 4:** Gene-Ontology (GO) functional analysis derived from proteomics.

**Supplementary Table 5:** Functional analysis of DEPs exclusively modulated by a SARS-CoV-2 structural protein.

**Supplementary Table 6:** Immunological functionality derived from the interlocking of transcriptome and proteome datasets.

## Author Contributions

Conceptualization, E.S.; methodology, M.L.-M; N.M.; K.A.; M.E.; E.B.; L.C.; M.dT; G.K.; D.E.; J. F.-I; E.S.; software, M.L.-M.; M.dT; J.F.-I; validation, N.M; formal analysis, M.L.-M.; E.S.; omics, K. A; J.F-I; M.dT.; investigation, M.L.-M; N.M.; M.E.; E.B.; L.C.; G.K.; D.E.; J.F.-I; E.S.; data curation, M.dT.; E.S.; writing—original draft preparation, E.S.; supervision, G.K.; D.E.; E.S.; funding acquisition, D.E.; J.F.-I; E.S. All authors have read and agreed to the published version of the manuscript.

## Data Availability Statement

Transcriptomic and MS data and results files were deposited in GEO accession GSE182849 (To review, https://www.ncbi.nlm.nih.gov/geo/query/acc.cgi?acc=GSE182849; Password: gdejkmiwznsxxgl) and ProteomeXchange Consortium via the JPOST partner with the identifier PXD027645 for Pro-teomeXchange and JPST001274 for jPOST (for reviewers: https://repository.jpostdb.org/preview/6417285016102b4b16aaa0; Access key: 3355).

## Acknowledgments

Authors thank all JPOST Team for helping with the mass spectrometric data deposit in ProteomeXChange/PRIDE. The Proteomics Platform of Navarrabiomed was member of Proteored (PRB3-ISCIII) supported by grant PT17/0019/009, of the PE I+D+I 2013-2016 funded by ISCIII and FEDER. The Clinical Neuroproteomics Unit of Navarrabiomed is member of the Global Consortium for Chemosensory Research (GCCR) and the Spanish Olfactory Network (ROE) (supported by grant RED2018-102662-T funded by Spanish Ministry of Science and Innovation).

## References

Appelberg, S., Gupta, S., Svensson Akusjarvi, S., Ambikan, A.T., Mikaeloff, F., Saccon, E., Vegvari, A., Benfeitas, R., Sperk, M., Stahlberg, M., Krishnan, S., Singh, K., Penninger, J.M., Mirazimi, A., and Neogi, U. (2020). Dysregulation in Akt/mTOR/HIF-1 signaling identified by proteo-transcriptomics of SARS-CoV-2 infected cells. Emerg Microbes Infect 9, 1748–1760.

Artal-Sanz, M., and Tavernarakis, N. (2009). Prohibitin and mitochondrial biology. Trends Endocrinol Metab 20, 394–401.

Bader, G.D., and Hogue, C.W. (2003). An automated method for finding molecular complexes in large protein interaction networks. BMC Bioinformatics 4, 2.

Bagheri, S.H., Asghari, A., Farhadi, M., Shamshiri, A.R., Kabir, A., Kamrava, S.K., Jalessi, M., Mohebbi, A., Alizadeh, R., Honarmand, A.A., Ghalehbaghi, B., Salimi, A., and Dehghani Firouzabadi, F. (2020). Coincidence of COVID-19 epidemic and olfactory dysfunction outbreak in Iran. Med J Islam Repub Iran 34, 62.

Beltran-Corbellini, A., Chico-Garcia, J.L., Martinez-Poles, J., Rodriguez-Jorge, F., Natera-Villalba, E., Gomez-Corral, J., Gomez-Lopez, A., Monreal, E., Parra-Diaz, P., Cortes-Cuevas, J.L., Galan, J.C., Fragola-Arnau, C., Porta-Etessam, J., Masjuan, J., and Alonso-Canovas, A. (2020). Acute-onset smell and taste disorders in the context of COVID-19: a pilot multicentre polymerase chain reaction based case-control study. Eur J Neurol 27, 1738–1741.

Biering, S.B., De Sousa, F.T.G., Tjang, L.V., Pahmeier, F., Ruan, R., Blanc, S.F., Patel, T.S., Worthington, C.M., Glasner, D.R., Castillo-Rojas, B., Servellita, V., Lo, N.T.N., Wong, M.P., Warnes, C.M., Sandoval, D.R., Clausen, T.M., Santos, Y.A., Ortega, V., Aguilar, H.C., Esko, J.D., Chui, C.Y., Pak, J.E., Beatty, P.R., and Harris, E. (2021). SARS-CoV-2 Spike triggers barrier dysfunction and vascular leak via integrins and TGF-beta signaling. bioRxiv.

Boson, B., Legros, V., Zhou, B., Siret, E., Mathieu, C., Cosset, F.L., Lavillette, D., and Denolly, S. (2021). The SARS-CoV-2 envelope and membrane proteins modulate maturation and retention of the spike protein, allowing assembly of virus-like particles. J Biol Chem 296, 100111.

Bouhaddou, M., Memon, D., Meyer, B., White, K.M., Rezelj, V.V., Correa Marrero, M., Polacco, B.J., Melnyk, J.E., Ulferts, S., Kaake, R.M., Batra, J., Richards, A.L., Stevenson, E., Gordon, D.E., Rojc, A., Obernier, K., Fabius, J.M., Soucheray, M., Miorin, L., Moreno, E., Koh, C., Tran, Q.D., Hardy, A., Robinot, R., Vallet, T., Nilsson-Payant, B.E., Hernandez-Armenta, C., Dunham, A., Weigang, S., Knerr, J., Modak, M., Quintero, D., Zhou, Y., Dugourd, A., Valdeolivas, A., Patil, T., Li, Q., Huttenhain, R., Cakir, M., Muralidharan, M., Kim, M., Jang, G., Tutuncuoglu, B., Hiatt, J., Guo, J.Z., Xu, J., Bouhaddou, S., Mathy, C.J.P., Gaulton, A., Manners, E.J., Felix, E., Shi, Y., Goff, M., Lim, J.K., Mcbride, T., O’neal, M.C., Cai, Y., Chang, J.C.J., Broadhurst, D.J., Klippsten, S., De Wit, E., Leach, A.R., Kortemme, T., Shoichet, B., Ott, M., Saez-Rodriguez, J., Tenoever, B.R., Mullins, R.D., Fischer, E.R., Kochs, G., Grosse, R., Garcia-Sastre, A., Vignuzzi, M., Johnson, J.R., Shokat, K.M., Swaney, D.L., Beltrao, P., and Krogan, N.J. (2020). The Global Phosphorylation Landscape of SARS-CoV-2 Infection. Cell 182, 685–712 e619.

Brann, D.H., Tsukahara, T., Weinreb, C., Lipovsek, M., Van Den Berge, K., Gong, B., Chance, R., Macaulay, I.C., Chou, H.J., Fletcher, R.B., Das, D., Street, K., De Bezieux, H.R., Choi, Y.G., Risso, D., Dudoit, S., Purdom, E., Mill, J., Hachem, R.A., Matsunami, H., Logan, D.W., Goldstein, B.J., Grubb, M.S., Ngai, J., and Datta, S.R. (2020). Non-neuronal expression of SARS-CoV-2 entry genes in the olfactory system suggests mechanisms underlying COVID-19-associated anosmia. Sci Adv 6.

Bryche, B., Le Bourhis, M., Congar, P., Martin, C., Rampin, O., and Meunier, N. (2019). Endothelin impacts on olfactory processing in rats. Behav Brain Res 362, 1–6.

Butowt, R., Meunier, N., Bryche, B., and Von Bartheld, C.S. (2021). The olfactory nerve is not a likely route to brain infection in COVID-19: a critical review of data from humans and animal models. Acta Neuropathol 141, 809–822.

Cheng, Y., Sun, F., Wang, L., Gao, M., Xie, Y., Sun, Y., Liu, H., Yuan, Y., Yi, W., Huang, Z., Yan, H., Peng, K., Wu, Y., and Cao, Z. (2020). Virus-induced p38 MAPK activation facilitates viral infection. Theranostics 10, 12223–12240.

Cooper, K.W., Brann, D.H., Farruggia, M.C., Bhutani, S., Pellegrino, R., Tsukahara, T., Weinreb, C., Joseph, P.V., Larson, E.D., Parma, V., Albers, M.W., Barlow, L.A., Datta, S.R., and Di Pizio, A. (2020). COVID-19 and the Chemical Senses: Supporting Players Take Center Stage. Neuron 107, 219–233.

De Haan, C.A., and Rottier, P.J. (2005). Molecular interactions in the assembly of coronaviruses. Adv Virus Res 64, 165–230.

De Melo, G.D., Lazarini, F., Levallois, S., Hautefort, C., Michel, V., Larrous, F., Verillaud, B., Aparicio, C., Wagner, S., Gheusi, G., Kergoat, L., Kornobis, E., Donati, F., Cokelaer, T., Hervochon, R., Madec, Y., Roze, E., Salmon, D., Bourhy, H., Lecuit, M., and Lledo, P.M. (2021). COVID-19-related anosmia is associated with viral persistence and inflammation in human olfactory epithelium and brain infection in hamsters. Sci Transl Med 13.

Deigendesch, N., Sironi, L., Kutza, M., Wischnewski, S., Fuchs, V., Hench, J., Frank, A., Nienhold, R., Mertz, K.D., Cathomas, G., Matter, M.S., Siegemund, M., Tolnay, M., Schirmer, L., Probstel, A.K., Tzankov, A., and Frank, S. (2020). Correlates of critical illness-related encephalopathy predominate postmortem COVID-19 neuropathology. Acta Neuropathol 140, 583–586.

Duart, G., Garcia-Murria, M.J., Grau, B., Acosta-Caceres, J.M., Martinez-Gil, L., and Mingarro, I. (2020). SARS-CoV-2 envelope protein topology in eukaryotic membranes. Open Biol 10, 200209.

Dube, M., Le Coupanec, A., Wong, A.H.M., Rini, J.M., Desforges, M., and Talbot, P.J. (2018). Axonal Transport Enables Neuron-to-Neuron Propagation of Human Coronavirus OC43. J Virol 92.

Durrant, D.M., Ghosh, S., and Klein, R.S. (2016). The Olfactory Bulb: An Immunosensory Effector Organ during Neurotropic Viral Infections. ACS Chem Neurosci 7, 464–469.

Ebrahim, A., Brunk, E., Tan, J., O’brien, E.J., Kim, D., Szubin, R., Lerman, J.A., Lechner, A., Sastry, A., Bordbar, A., Feist, A.M., and Palsson, B.O. (2016). Multi-omic data integration enables discovery of hidden biological regularities. Nat Commun 7, 13091.

Escors, D., Lopes, L., Lin, R., Hiscott, J., Akira, S., Davis, R.J., and Collins, M.K. (2008). Targeting dendritic cell signaling to regulate the response to immunization. Blood 111, 3050–3061.

Escors, D., Ortego, J., Laude, H., and Enjuanes, L. (2001). The membrane M protein carboxy terminus binds to transmissible gastroenteritis coronavirus core and contributes to core stability. J Virol 75, 1312–1324.

Fagone, P., Ciurleo, R., Lombardo, S.D., Iacobello, C., Palermo, C.I., Shoenfeld, Y., Bendtzen, K., Bramanti, P., and Nicoletti, F. (2020). Transcriptional landscape of SARS-CoV-2 infection dismantles pathogenic pathways activated by the virus, proposes unique sex-specific differences and predicts tailored therapeutic strategies. Autoimmun Rev 19, 102571.

Flynn, R.A., Belk, J.A., Qi, Y., Yasumoto, Y., Wei, J., Alfajaro, M.M., Shi, Q., Mumbach, M.R., Limaye, A., Deweirdt, P.C., Schmitz, C.O., Parker, K.R., Woo, E., Chang, H.Y., Horvath, T.L., Carette, J.E., Bertozzi, C.R., Wilen, C.B., and Satpathy, A.T. (2021). Discovery and functional interrogation of SARS-CoV-2 RNA-host protein interactions. Cell 184, 2394–2411 e2316.

Fung, T.S., and Liu, D.X. (2017). Activation of the c-Jun NH2-terminal kinase pathway by coronavirus infectious bronchitis virus promotes apoptosis independently of c-Jun. Cell Death Dis 8, 3215.

Gato-Canas, M., Zuazo, M., Arasanz, H., Ibanez-Vea, M., Lorenzo, L., Fernandez-Hinojal, G., Vera, R., Smerdou, C., Martisova, E., Arozarena, I., Wellbrock, C., Llopiz, D., Ruiz, M., Sarobe, P., Breckpot, K., Kochan, G., and Escors, D. (2017). PDL1 Signals through Conserved Sequence Motifs to Overcome Interferon-Mediated Cytotoxicity. Cell Rep 20, 1818–1829.

Gerkin, R.C., Ohla, K., Veldhuizen, M.G., Joseph, P.V., Kelly, C.E., Bakke, A.J., Steele, K.E., Farruggia, M.C., Pellegrino, R., Pepino, M.Y., Bouysset, C., Soler, G.M., Pereda-Loth, V., Dibattista, M., Cooper, K.W., Croijmans, I., Di Pizio, A., Ozdener, M.H., Fjaeldstad, A.W., Lin, C., Sandell, M.A., Singh, P.B., Brindha, V.E., Olsson, S.B., Saraiva, L.R., Ahuja, G., Alwashahi, M.K., Bhutani, S., D’errico, A., Fornazieri, M.A., Golebiowski, J., Dar Hwang, L., Ozturk, L., Roura, E., Spinelli, S., Whitcroft, K.L., Faraji, F., Fischmeister, F.P.S., Heinbockel, T., Hsieh, J.W., Huart, C., Konstantinidis, I., Menini, A., Morini, G., Olofsson, J.K., Philpott, C.M., Pierron, D., Shields, V.D.C., Voznessenskaya, V.V., Albayay, J., Altundag, A., Bensafi, M., Bock, M.A., Calcinoni, O., Fredborg, W., Laudamiel, C., Lim, J., Lundstrom, J.N., Macchi, A., Meyer, P., Moein, S.T., Santamaria, E., Sengupta, D., Rohlfs Dominguez, P., Yanik, H., Hummel, T., Hayes, J.E., Reed, D.R., Niv, M.Y., Munger, S.D., Parma, V., and Author, G.G. (2021). Recent Smell Loss Is the Best Predictor of COVID-19 Among Individuals With Recent Respiratory Symptoms. Chem Senses 46.

Giacomelli, A., Pezzati, L., Conti, F., Bernacchia, D., Siano, M., Oreni, L., Rusconi, S., Gervasoni, C., Ridolfo, A.L., Rizzardini, G., Antinori, S., and Galli, M. (2020). Self-reported Olfactory and Taste Disorders in Patients With Severe Acute Respiratory Coronavirus 2 Infection: A Cross-sectional Study. Clin Infect Dis 71, 889–890.

Gibellini, L., De Biasi, S., Paolini, A., Borella, R., Boraldi, F., Mattioli, M., Lo Tartaro, D., Fidanza, L., Caro-Maldonado, A., Meschiari, M., Iadisernia, V., Bacca, E., Riva, G., Cicchetti, L., Quaglino, D., Guaraldi, G., Busani, S., Girardis, M., Mussini, C., and Cossarizza, A. (2020). Altered bioenergetics and mitochondrial dysfunction of monocytes in patients with COVID-19 pneumonia. EMBO Mol Med 12, e13001.

Gillet, L.C., Navarro, P., Tate, S., Rost, H., Selevsek, N., Reiter, L., Bonner, R., and Aebersold, R. (2012). Targeted data extraction of the MS/MS spectra generated by data-independent acquisition: a new concept for consistent and accurate proteome analysis. Mol Cell Proteomics 11, O111 016717.

Grimes, J.M., and Grimes, K.V. (2020). p38 MAPK inhibition: A promising therapeutic approach for COVID-19. J Mol Cell Cardiol 144, 63–65.

Guzik, T.J., Mohiddin, S.A., Dimarco, A., Patel, V., Savvatis, K., Marelli-Berg, F.M., Madhur, M.S., Tomaszewski, M., Maffia, P., D’acquisto, F., Nicklin, S.A., Marian, A.J., Nosalski, R., Murray, E.C., Guzik, B., Berry, C., Touyz, R.M., Kreutz, R., Wang, D.W., Bhella, D., Sagliocco, O., Crea, F., Thomson, E.C., and Mcinnes, I.B. (2020). COVID-19 and the cardiovascular system: implications for risk assessment, diagnosis, and treatment options. Cardiovasc Res 116, 1666–1687.

Haehner, A., Draf, J., Drager, S., De With, K., and Hummel, T. (2020). Predictive Value of Sudden Olfactory Loss in the Diagnosis of COVID-19. ORL J Otorhinolaryngol Relat Spec 82, 175–180.

Hatton, C.F., Botting, R.A., Duenas, M.E., Haq, I.J., Verdon, B., Thompson, B.J., Spegarova, J.S., Gothe, F., Stephenson, E., Gardner, A.I., Murphy, S., Scott, J., Garnett, J.P., Carrie, S., Powell, J., Khan, C.M.A., Huang, L., Hussain, R., Coxhead, J., Davey, T., Simpson, A.J., Haniffa, M., Hambleton, S., Brodlie, M., Ward, C., Trost, M., Reynolds, G., and Duncan, C.J.A. (2021). Delayed induction of type I and III interferons mediates nasal epithelial cell permissiveness to SARS-CoV-2. Nat Commun 12, 7092.

Huang, C., Wang, Y., Li, X., Ren, L., Zhao, J., Hu, Y., Zhang, L., Fan, G., Xu, J., Gu, X., Cheng, Z., Yu, T., Xia, J., Wei, Y., Wu, W., Xie, X., Yin, W., Li, H., Liu, M., Xiao, Y., Gao, H., Guo, L., Xie, J., Wang, G., Jiang, R., Gao, Z., Jin, Q., Wang, J., and Cao, B. (2020). Clinical features of patients infected with 2019 novel coronavirus in Wuhan, China. Lancet 395, 497–506.

Hue, S., Beldi-Ferchiou, A., Bendib, I., Surenaud, M., Fourati, S., Frapard, T., Rivoal, S., Razazi, K., Carteaux, G., Delfau-Larue, M.H., Mekontso-Dessap, A., Audureau, E., and De Prost, N. (2020). Uncontrolled Innate and Impaired Adaptive Immune Responses in Patients with COVID-19 Acute Respiratory Distress Syndrome. Am J Respir Crit Care Med 202, 1509–1519.

Huntington, K.E., Carlsen, L., So, E.Y., Piesche, M., Liang, O., and El-Deiry, W.S. (2022). Integrin/TGF-beta1 inhibitor GLPG-0187 blocks SARS-CoV-2 Delta and Omicron pseudovirus infection of airway epithelial cells which could attenuate disease severity. medRxiv.

Jensen, E.C. (2012). Use of fluorescent probes: their effect on cell biology and limitations. Anat Rec (Hoboken) 295, 2031–2036.

Jiao, L., Yang, Y., Yu, W., Zhao, Y., Long, H., Gao, J., Ding, K., Ma, C., Li, J., Zhao, S., Wang, H., Li, H., Yang, M., Xu, J., Wang, J., Yang, J., Kuang, D., Luo, F., Qian, X., Xu, L., Yin, B., Liu, W., Liu, H., Lu, S., and Peng, X. (2021). The olfactory route is a potential way for SARS-CoV-2 to invade the central nervous system of rhesus monkeys. Signal Transduct Target Ther 6, 169.

Kim, D., Paggi, J.M., Park, C., Bennett, C., and Salzberg, S.L. (2019). Graph-based genome alignment and genotyping with HISAT2 and HISAT-genotype. Nat Biotechnol 37, 907–915.

Kindrachuk, J., Ork, B., Hart, B.J., Mazur, S., Holbrook, M.R., Frieman, M.B., Traynor, D., Johnson, R.F., Dyall, J., Kuhn, J.H., Olinger, G.G., Hensley, L.E., and Jahrling, P.B. (2015). Antiviral potential of ERK/MAPK and PI3K/AKT/mTOR signaling modulation for Middle East respiratory syndrome coronavirus infection as identified by temporal kinome analysis. Antimicrob Agents Chemother 59, 1088–1099.

Kopylova, E., Noe, L., and Touzet, H. (2012). SortMeRNA: fast and accurate filtering of ribosomal RNAs in metatranscriptomic data. Bioinformatics 28, 3211–3217.

Lachen-Montes, M., Corrales, F.J., Fernandez-Irigoyen, J., and Santamaria, E. (2020). Proteomics Insights Into the Molecular Basis of SARS-CoV-2 Infection: What We Can Learn From the Human Olfactory Axis. Front Microbiol 11, 2101.

Lachen-Montes, M., Gonzalez-Morales, A., Palomino, M., Ausin, K., Gomez-Ochoa, M., Zelaya, M.V., Ferrer, I., Perez-Mediavilla, A., Fernandez-Irigoyen, J., and Santamaria, E. (2019). Early-Onset Molecular Derangements in the Olfactory Bulb of Tg2576 Mice: Novel Insights Into the Stress-Responsive Olfactory Kinase Dynamics in Alzheimer’s Disease. Front Aging Neurosci 11, 141.

Lanning, N.J., Looyenga, B.D., Kauffman, A.L., Niemi, N.M., Sudderth, J., Deberardinis, R.J., and Mackeigan, J.P. (2014). A mitochondrial RNAi screen defines cellular bioenergetic determinants and identifies an adenylate kinase as a key regulator of ATP levels. Cell Rep 7, 907–917.

Le Bourhis, M., Rimbaud, S., Grebert, D., Congar, P., and Meunier, N. (2014). Endothelin uncouples gap junctions in sustentacular cells and olfactory ensheathing cells of the olfactory mucosa. Eur J Neurosci 40, 2878–2887.

Li, H., Handsaker, B., Wysoker, A., Fennell, T., Ruan, J., Homer, N., Marth, G., Abecasis, G., Durbin, R., and Genome Project Data Processing, S. (2009). The Sequence Alignment/Map format and SAMtools. Bioinformatics 25, 2078–2079.

Li, Q., Guan, X., Wu, P., Wang, X., Zhou, L., Tong, Y., Ren, R., Leung, K.S.M., Lau, E.H.Y., Wong, J.Y., Xing, X., Xiang, N., Wu, Y., Li, C., Chen, Q., Li, D., Liu, T., Zhao, J., Liu, M., Tu, W., Chen, C., Jin, L., Yang, R., Wang, Q., Zhou, S., Wang, R., Liu, H., Luo, Y., Liu, Y., Shao, G., Li, H., Tao, Z., Yang, Y., Deng, Z., Liu, B., Ma, Z., Zhang, Y., Shi, G., Lam, T.T.Y., Wu, J.T., Gao, G.F., Cowling, B.J., Yang, B., Leung, G.M., and Feng, Z. (2020). Early Transmission Dynamics in Wuhan, China, of Novel Coronavirus-Infected Pneumonia. N Engl J Med 382, 1199–1207.

Liao, Y., Smyth, G.K., and Shi, W. (2014). featureCounts: an efficient general purpose program for assigning sequence reads to genomic features. Bioinformatics 30, 923–930.

Liu, R., Strom, A.L., Zhai, J., Gal, J., Bao, S., Gong, W., and Zhu, H. (2009). Enzymatically inactive adenylate kinase 4 interacts with mitochondrial ADP/ATP translocase. Int J Biochem Cell Biol 41, 1371–1380.

Liu, X., Verma, A., Garcia, G., Ramage, H., Myers, R.L., Lucas, A., Michaelson, J.J., Coryell, W., Kumar, A., Charney, A.W., Kazanietz, M.G., Rader, D.J., Ritchie, M.D., Berrettini, W.H., Schultz, D.C., Cherry, S., Damoiseaux, R., Arumugaswami, V., and Klein, P.S. (2021). Targeting the Coronavirus Nucleocapsid Protein through GSK-3 Inhibition. medRxiv.

Lopez, G., Tonello, C., Osipova, G., Carsana, L., Biasin, M., Cappelletti, G., Pellegrinelli, A., Lauri, E., Zerbi, P., Rossi, R.S., and Nebuloni, M. (2021). Olfactory bulb SARS-CoV-2 infection is not paralleled by the presence of virus in other central nervous system areas. Neuropathol Appl Neurobiol.

Lou, J.J., Movassaghi, M., Gordy, D., Olson, M.G., Zhang, T., Khurana, M.S., Chen, Z., Perez-Rosendahl, M., Thammachantha, S., Singer, E.J., Magaki, S.D., Vinters, H.V., and Yong, W.H. (2021). Neuropathology of COVID-19 (neuro-COVID): clinicopathological update. Free Neuropathol 2.

Love, M.I., Huber, W., and Anders, S. (2014). Moderated estimation of fold change and dispersion for RNA-seq data with DESeq2. Genome Biol 15, 550.

Mahmud, F.J., Du, Y., Greif, E., Boucher, T., Dannals, R.F., Mathews, W.B., Pomper, M.G., Sysa-Shah, P., Metcalf Pate, K.A., Lyons, C., Carlson, B., Chacona, M., and Brown, A.M. (2020). Osteopontin/secreted phosphoprotein-1 behaves as a molecular brake regulating the neuroinflammatory response to chronic viral infection. J Neuroinflammation 17, 273.

Mann, M., Kumar, C., Zeng, W.F., and Strauss, M.T. (2021). Artificial intelligence for proteomics and biomarker discovery. Cell Syst 12, 759–770.

Matschke, J., Lutgehetmann, M., Hagel, C., Sperhake, J.P., Schroder, A.S., Edler, C., Mushumba, H., Fitzek, A., Allweiss, L., Dandri, M., Dottermusch, M., Heinemann, A., Pfefferle, S., Schwabenland, M., Sumner Magruder, D., Bonn, S., Prinz, M., Gerloff, C., Puschel, K., Krasemann, S., Aepfelbacher, M., and Glatzel, M. (2020). Neuropathology of patients with COVID-19 in Germany: a post-mortem case series. Lancet Neurol 19, 919–929.

Meinhardt, J., Radke, J., Dittmayer, C., Franz, J., Thomas, C., Mothes, R., Laue, M., Schneider, J., Brunink, S., Greuel, S., Lehmann, M., Hassan, O., Aschman, T., Schumann, E., Chua, R.L., Conrad, C., Eils, R., Stenzel, W., Windgassen, M., Rossler, L., Goebel, H.H., Gelderblom, H.R., Martin, H., Nitsche, A., Schulz-Schaeffer, W.J., Hakroush, S., Winkler, M.S., Tampe, B., Scheibe, F., Kortvelyessy, P., Reinhold, D., Siegmund, B., Kuhl, A.A., Elezkurtaj, S., Horst, D., Oesterhelweg, L., Tsokos, M., Ingold-Heppner, B., Stadelmann, C., Drosten, C., Corman, V.M., Radbruch, H., and Heppner, F.L. (2021). Olfactory transmucosal SARS-CoV-2 invasion as a port of central nervous system entry in individuals with COVID-19. Nat Neurosci 24, 168–175.

Menni, C., Valdes, A.M., Freidin, M.B., Sudre, C.H., Nguyen, L.H., Drew, D.A., Ganesh, S., Varsavsky, T., Cardoso, M.J., El-Sayed Moustafa, J.S., Visconti, A., Hysi, P., Bowyer, R.C.E., Mangino, M., Falchi, M., Wolf, J., Ourselin, S., Chan, A.T., Steves, C.J., and Spector, T.D. (2020). Real-time tracking of self-reported symptoms to predict potential COVID-19. Nat Med 26, 1037–1040.

Menter, T., Haslbauer, J.D., Nienhold, R., Savic, S., Hopfer, H., Deigendesch, N., Frank, S., Turek, D., Willi, N., Pargger, H., Bassetti, S., Leuppi, J.D., Cathomas, G., Tolnay, M., Mertz, K.D., and Tzankov, A. (2020). Postmortem examination of COVID-19 patients reveals diffuse alveolar damage with severe capillary congestion and variegated findings in lungs and other organs suggesting vascular dysfunction. Histopathology 77, 198–209.

Merkwirth, C., Martinelli, P., Korwitz, A., Morbin, M., Bronneke, H.S., Jordan, S.D., Rugarli, E.I., and Langer, T. (2012). Loss of prohibitin membrane scaffolds impairs mitochondrial architecture and leads to tau hyperphosphorylation and neurodegeneration. PLoS Genet 8, e1003021.

Morbini, P., Benazzo, M., Verga, L., Pagella, F.G., Mojoli, F., Bruno, R., and Marena, C. (2020). Ultrastructural Evidence of Direct Viral Damage to the Olfactory Complex in Patients Testing Positive for SARS-CoV-2. JAMA Otolaryngol Head Neck Surg 146, 972–973.

Naqvi, A.a.T., Fatima, K., Mohammad, T., Fatima, U., Singh, I.K., Singh, A., Atif, S.M., Hariprasad, G., Hasan, G.M., and Hassan, M.I. (2020). Insights into SARS-CoV-2 genome, structure, evolution, pathogenesis and therapies: Structural genomics approach. Biochim Biophys Acta Mol Basis Dis 1866, 165878.

Nie, X., Qian, L., Sun, R., Huang, B., Dong, X., Xiao, Q., Zhang, Q., Lu, T., Yue, L., Chen, S., Li, X., Sun, Y., Li, L., Xu, L., Li, Y., Yang, M., Xue, Z., Liang, S., Ding, X., Yuan, C., Peng, L., Liu, W., Yi, X., Lyu, M., Xiao, G., Xu, X., Ge, W., He, J., Fan, J., Wu, J., Luo, M., Chang, X., Pan, H., Cai, X., Zhou, J., Yu, J., Gao, H., Xie, M., Wang, S., Ruan, G., Chen, H., Su, H., Mei, H., Luo, D., Zhao, D., Xu, F., Li, Y., Zhu, Y., Xia, J., Hu, Y., and Guo, T. (2021). Multi-organ proteomic landscape of COVID-19 autopsies. Cell 184, 775–791 e714.

Okonechnikov, K., Conesa, A., and Garcia-Alcalde, F. (2016). Qualimap 2: advanced multi-sample quality control for high-throughput sequencing data. Bioinformatics 32, 292–294.

Okuda, S., Watanabe, Y., Moriya, Y., Kawano, S., Yamamoto, T., Matsumoto, M., Takami, T., Kobayashi, D., Araki, N., Yoshizawa, A.C., Tabata, T., Sugiyama, N., Goto, S., and Ishihama, Y. (2017). jPOSTrepo: an international standard data repository for proteomes. Nucleic Acids Res 45, D1107–D1111.

Parma, V., Ohla, K., Veldhuizen, M.G., Niv, M.Y., Kelly, C.E., Bakke, A.J., Cooper, K.W., Bouysset, C., Pirastu, N., Dibattista, M., Kaur, R., Liuzza, M.T., Pepino, M.Y., Schopf, V., Pereda-Loth, V., Olsson, S.B., Gerkin, R.C., Rohlfs Dominguez, P., Albayay, J., Farruggia, M.C., Bhutani, S., Fjaeldstad, A.W., Kumar, R., Menini, A., Bensafi, M., Sandell, M., Konstantinidis, I., Di Pizio, A., Genovese, F., Ozturk, L., Thomas-Danguin, T., Frasnelli, J., Boesveldt, S., Saatci, O., Saraiva, L.R., Lin, C., Golebiowski, J., Hwang, L.D., Ozdener, M.H., Guardia, M.D., Laudamiel, C., Ritchie, M., Havlicek, J., Pierron, D., Roura, E., Navarro, M., Nolden, A.A., Lim, J., Whitcroft, K.L., Colquitt, L.R., Ferdenzi, C., Brindha, E.V., Altundag, A., Macchi, A., Nunez-Parra, A., Patel, Z.M., Fiorucci, S., Philpott, C.M., Smith, B.C., Lundstrom, J.N., Mucignat, C., Parker, J.K., Van Den Brink, M., Schmuker, M., Fischmeister, F.P.S., Heinbockel, T., Shields, V.D.C., Faraji, F., Santamaria, E., Fredborg, W.E.A., Morini, G., Olofsson, J.K., Jalessi, M., Karni, N., D’errico, A., Alizadeh, R., Pellegrino, R., Meyer, P., Huart, C., Chen, B., Soler, G.M., Alwashahi, M.K., Welge-Lussen, A., Freiherr, J., De Groot, J.H.B., Klein, H., Okamoto, M., Singh, P.B., Hsieh, J.W., Author, G.G., Reed, D.R., Hummel, T., Munger, S.D., and Hayes, J.E. (2020). More Than Smell-COVID-19 Is Associated With Severe Impairment of Smell, Taste, and Chemesthesis. Chem Senses 45, 609–622.

Pervushin, K., Tan, E., Parthasarathy, K., Lin, X., Jiang, F.L., Yu, D., Vararattanavech, A., Soong, T.W., Liu, D.X., and Torres, J. (2009). Structure and inhibition of the SARS coronavirus envelope protein ion channel. PLoS Pathog 5, e1000511.

Pierron, D., Pereda-Loth, V., Mantel, M., Moranges, M., Bignon, E., Alva, O., Kabous, J., Heiske, M., Pacalon, J., David, R., Dinnella, C., Spinelli, S., Monteleone, E., Farruggia, M.C., Cooper, K.W., Sell, E.A., Thomas-Danguin, T., Bakke, A.J., Parma, V., Hayes, J.E., Letellier, T., Ferdenzi, C., Golebiowski, J., and Bensafi, M. (2020). Smell and taste changes are early indicators of the COVID-19 pandemic and political decision effectiveness. Nat Commun 11, 5152.

Piras, I.S., Huentelman, M.J., Walker, J.E., Arce, R., Glass, M.J., Vargas, D., Sue, L.I., Intorcia, A.J., Nelson, C.M., Suszczewicz, K.E., Borja, C.L., Desforges, M., Deture, M., Dickson, D.W., Beach, T.G., and Serrano, G.E. (2021). Olfactory Bulb and Amygdala Gene Expression Changes in Subjects Dying with COVID-19. medRxiv.

Powell, M.A., Black, R.T., Smith, T.L., Reeves, T.M., and Phillips, L.L. (2019). Matrix Metalloproteinase 9 and Osteopontin Interact to Support Synaptogenesis in the Olfactory Bulb after Mild Traumatic Brain Injury. J Neurotrauma 36, 1615–1631.

Rana, A.K., Rahmatkar, S.N., Kumar, A., and Singh, D. (2021). Glycogen synthase kinase-3: A putative target to combat severe acute respiratory syndrome coronavirus 2 (SARS-CoV-2) pandemic. Cytokine Growth Factor Rev 58, 92–101.

Rhea, E.M., Logsdon, A.F., Hansen, K.M., Williams, L.M., Reed, M.J., Baumann, K.K., Holden, S.J., Raber, J., Banks, W.A., and Erickson, M.A. (2021). The S1 protein of SARS-CoV-2 crosses the blood-brain barrier in mice. Nat Neurosci 24, 368–378.

Ricci, D., Etna, M.P., Rizzo, F., Sandini, S., Severa, M., and Coccia, E.M. (2021). Innate Immune Response to SARS-CoV-2 Infection: From Cells to Soluble Mediators. Int J Mol Sci 22.

Rudd, C.E. (2020). GSK-3 Inhibition as a Therapeutic Approach Against SARs CoV2: Dual Benefit of Inhibiting Viral Replication While Potentiating the Immune Response. Front Immunol 11, 1638.

Saccon, E., Chen, X., Mikaeloff, F., Rodriguez, J.E., Szekely, L., Vinhas, B.S., Krishnan, S., Byrareddy, S.N., Frisan, T., Vegvari, A., Mirazimi, A., Neogi, U., and Gupta, S. (2021). Cell-type-resolved quantitative proteomics map of interferon response against SARS-CoV-2. iScience 24, 102420.

Satarker, S., and Nampoothiri, M. (2020). Structural Proteins in Severe Acute Respiratory Syndrome Coronavirus-2. Arch Med Res 51, 482–491.

Schoeman, D., and Fielding, B.C. (2019). Coronavirus envelope protein: current knowledge. Virol J 16, 69.

Schwabenland, M., Salie, H., Tanevski, J., Killmer, S., Lago, M.S., Schlaak, A.E., Mayer, L., Matschke, J., Puschel, K., Fitzek, A., Ondruschka, B., Mei, H.E., Boettler, T., Neumann-Haefelin, C., Hofmann, M., Breithaupt, A., Genc, N., Stadelmann, C., Saez-Rodriguez, J., Bronsert, P., Knobeloch, K.P., Blank, T., Thimme, R., Glatzel, M., Prinz, M., and Bengsch, B. (2021). Deep spatial profiling of human COVID-19 brains reveals neuroinflammation with distinct microanatomical microglia-T-cell interactions. Immunity 54, 1594–1610 e1511.

Serrano, G.E., Walker, J.E., Arce, R., Glass, M.J., Vargas, D., Sue, L.I., Intorcia, A.J., Nelson, C.M., Oliver, J., Papa, J., Russell, A., Suszczewicz, K.E., Borja, C.I., Belden, C., Goldfarb, D., Shprecher, D., Atri, A., Adler, C.H., Shill, H.A., Driver-Dunckley, E., Mehta, S.H., Readhead, B., Huentelman, M.J., Peters, J.L., Alevritis, E., Bimi, C., Mizgerd, J.P., Reiman, E.M., Montine, T.J., Desforges, M., Zehnder, J.L., Sahoo, M.K., Zhang, H., Solis, D., Pinsky, B.A., Deture, M., Dickson, D.W., and Beach, T.G. (2021). Mapping of SARS-CoV-2 Brain Invasion and Histopathology in COVID-19 Disease. medRxiv.

Shang, J., Wan, Y., Luo, C., Ye, G., Geng, Q., Auerbach, A., and Li, F. (2020). Cell entry mechanisms of SARS-CoV-2. Proc Natl Acad Sci U S A 117, 11727–11734.

Stadnytskyi, V., Bax, C.E., Bax, A., and Anfinrud, P. (2020). The airborne lifetime of small speech droplets and their potential importance in SARS-CoV-2 transmission. Proc Natl Acad Sci U S A 117, 11875–11877.

Stukalov, A., Girault, V., Grass, V., Karayel, O., Bergant, V., Urban, C., Haas, D.A., Huang, Y., Oubraham, L., Wang, A., Hamad, M.S., Piras, A., Hansen, F.M., Tanzer, M.C., Paron, I., Zinzula, L., Engleitner, T., Reinecke, M., Lavacca, T.M., Ehmann, R., Wolfel, R., Jores, J., Kuster, B., Protzer, U., Rad, R., Ziebuhr, J., Thiel, V., Scaturro, P., Mann, M., and Pichlmair, A. (2021). Multilevel proteomics reveals host perturbations by SARS-CoV-2 and SARS-CoV. Nature 594, 246–252.

Tang, W.H., Shilov, I.V., and Seymour, S.L. (2008). Nonlinear fitting method for determining local false discovery rates from decoy database searches. J Proteome Res 7, 3661–3667.

Tugaeva, K.V., Hawkins, D., Smith, J.L.R., Bayfield, O.W., Ker, D.S., Sysoev, A.A., Klychnikov, O.I., Antson, A.A., and Sluchanko, N.N. (2021). The Mechanism of SARS-CoV-2 Nucleocapsid Protein Recognition by the Human 14-3-3 Proteins. J Mol Biol 433, 166875.

Tyanova, S., Temu, T., Sinitcyn, P., Carlson, A., Hein, M.Y., Geiger, T., Mann, M., and Cox, J. (2016). The Perseus computational platform for comprehensive analysis of (prote)omics data. Nat Methods 13, 731–740.

Varet, H., Brillet-Gueguen, L., Coppee, J.Y., and Dillies, M.A. (2016). SARTools: A DESeq2- and EdgeR-Based R Pipeline for Comprehensive Differential Analysis of RNA-Seq Data. PLoS One 11, e0157022.

Wang, W., Chen, J., Hu, D., Pan, P., Liang, L., Wu, W., Tang, Y., Huang, X.R., Yu, X., Wu, J., and Lan, H.Y. (2022). SARS-CoV-2 N Protein Induces Acute Kidney Injury via Smad3-Dependent G1 Cell Cycle Arrest Mechanism. Adv Sci (Weinh) 9, e2103248.

Willems, L.H., Nagy, M., Ten Cate, H., Spronk, H.M.H., Groh, L.A., Leentjens, J., Janssen, N.a.F., Netea, M.G., Thijssen, D.H.J., Hannink, G., Van Petersen, A.S., and Warle, M.C. (2021). Sustained inflammation, coagulation activation and elevated endothelin-1 levels without macrovascular dysfunction at 3 months after COVID-19. Thromb Res 209, 106–114.

Xu, W., Pei, G., Liu, H., Ju, X., Wang, J., Ding, Q., and Li, P. (2021). Compartmentalization-aided interaction screening reveals extensive high-order complexes within the SARS-CoV-2 proteome. Cell Rep 37, 109778.

Xydakis, M.S., Albers, M.W., Holbrook, E.H., Lyon, D.M., Shih, R.Y., Frasnelli, J.A., Pagenstecher, A., Kupke, A., Enquist, L.W., and Perlman, S. (2021). Post-viral effects of COVID-19 in the olfactory system and their implications. Lancet Neurol 20, 753–761.

Yang, J., Zheng, Y., Gou, X., Pu, K., Chen, Z., Guo, Q., Ji, R., Wang, H., Wang, Y., and Zhou, Y. (2020). Prevalence of comorbidities and its effects in patients infected with SARS-CoV-2: a systematic review and meta-analysis. Int J Infect Dis 94, 91–95.

Ye, Q., Zhou, J., He, Q., Li, R.T., Yang, G., Zhang, Y., Wu, S.J., Chen, Q., Shi, J.H., Zhang, R.R., Zhu, H.M., Qiu, H.Y., Zhang, T., Deng, Y.Q., Li, X.F., Liu, J.F., Xu, P., Yang, X., and Qin, C.F. (2021). SARS-CoV-2 infection in the mouse olfactory system. Cell Discov 7, 49.

Zhou, P., Qian, L., D’aurelio, M., Cho, S., Wang, G., Manfredi, G., Pickel, V., and Iadecola, C. (2012). Prohibitin reduces mitochondrial free radical production and protects brain cells from different injury modalities. J Neurosci 32, 583–592.

Zhou, Y., Zhou, B., Pache, L., Chang, M., Khodabakhshi, A.H., Tanaseichuk, O., Benner, C., and Chanda, S.K. (2019). Metascape provides a biologist-oriented resource for the analysis of systems-level datasets. Nat Commun 10, 1523.

