## Supplemental figures for "Metabolic dyshomeostasis induced by SARS-CoV-2 structural proteins reveals immunological insights into viral olfactory interactions"

### Slide 1
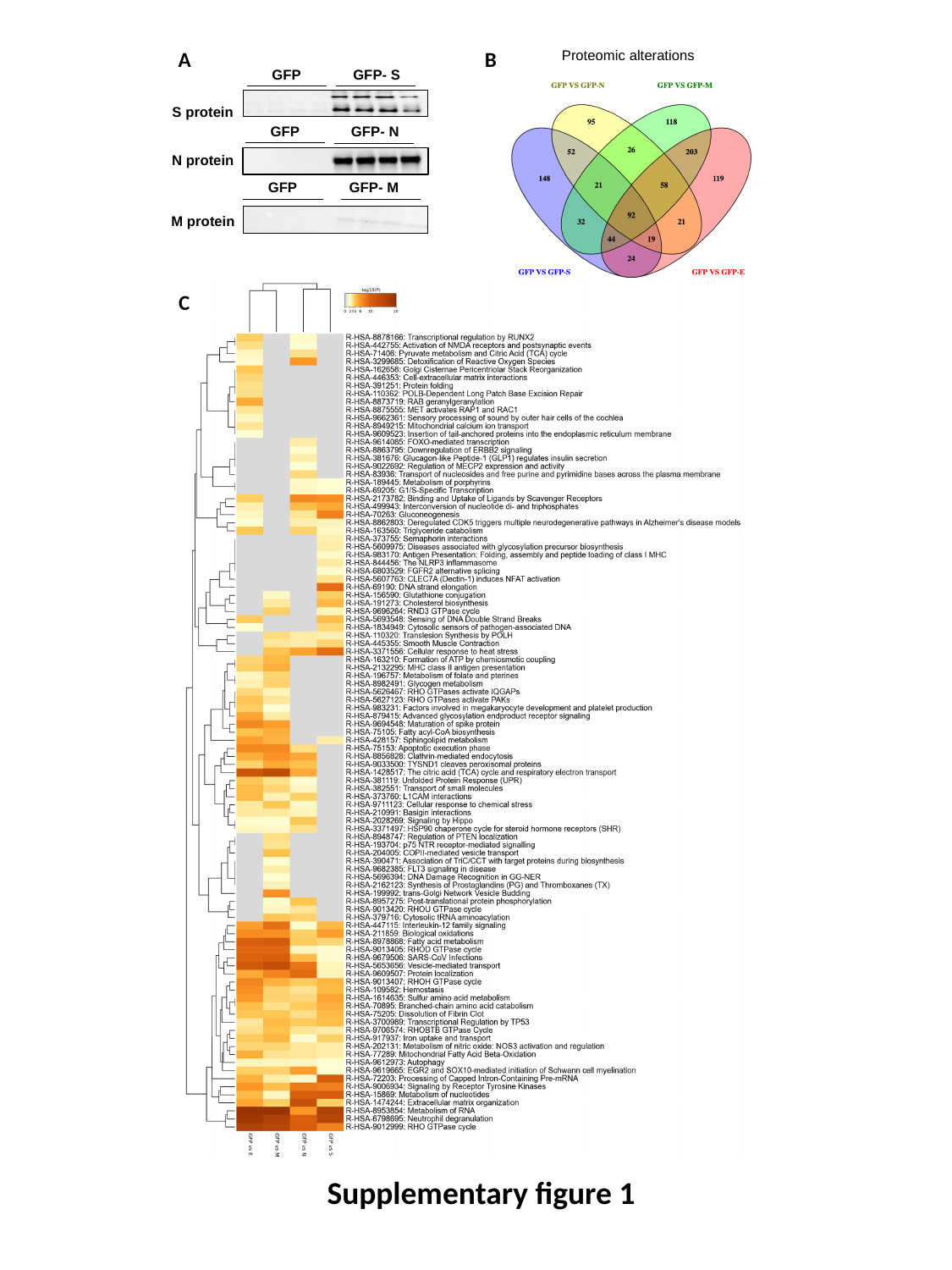

A
B
Proteomic alterations
GFP
GFP- S
S protein
GFP
GFP- N
N protein
GFP
GFP- M
M protein
C
Supplementary figure 1

### Slide 2
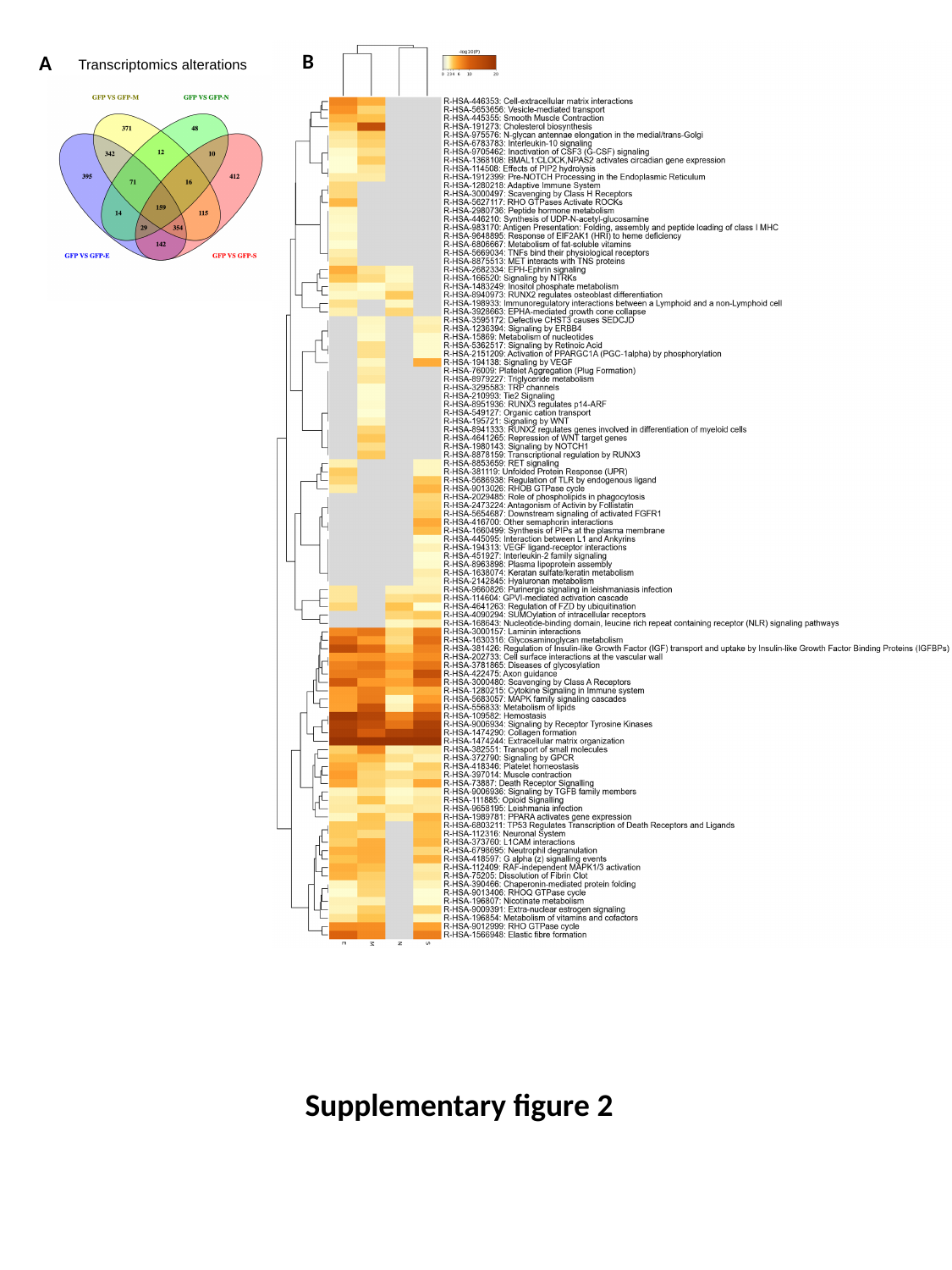

B
A
Transcriptomics alterations
Supplementary figure 2

### Slide 3
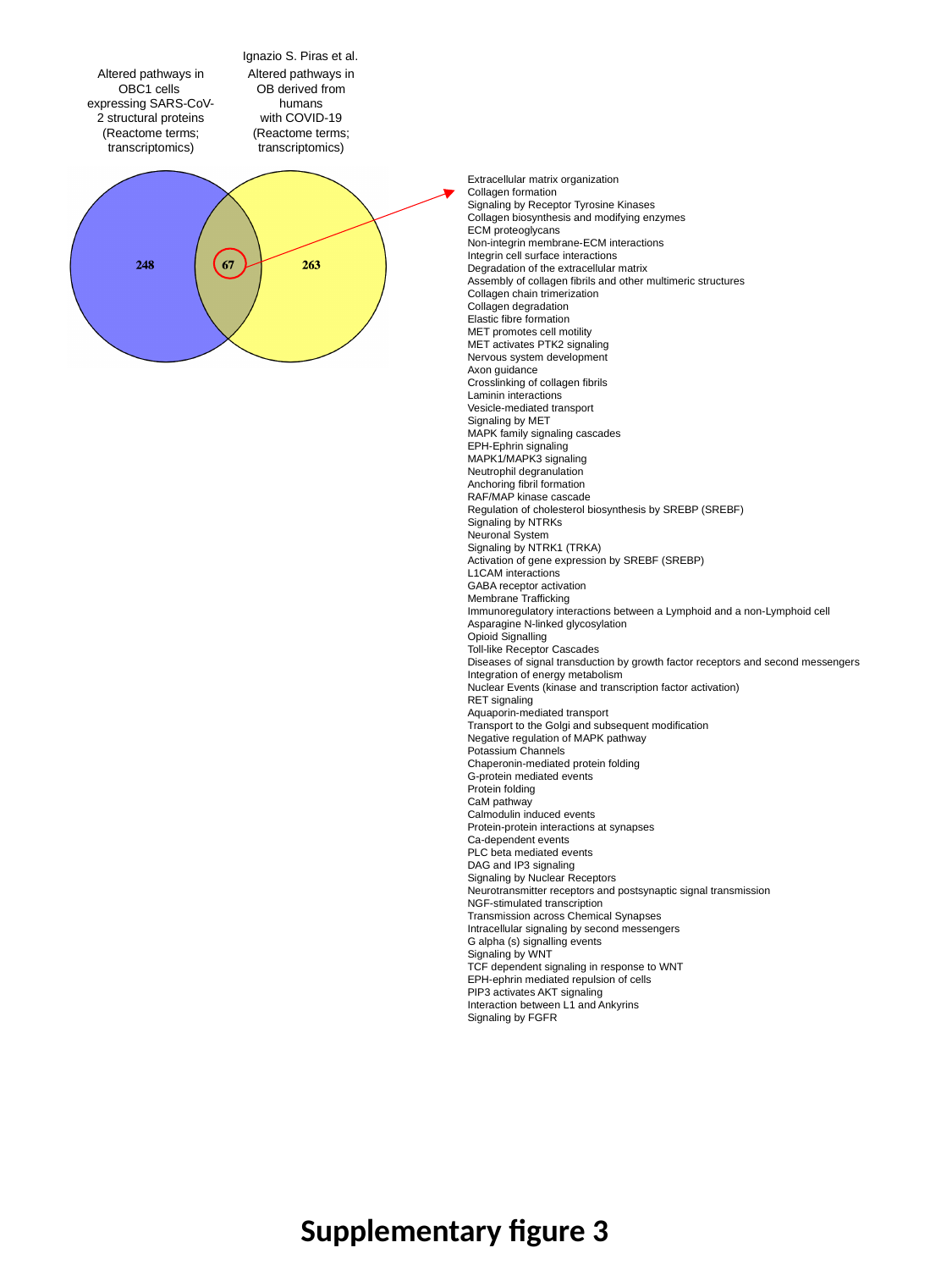

Ignazio S. Piras et al.
Altered pathways in OBC1 cells
expressing SARS-CoV-2 structural proteins
(Reactome terms; transcriptomics)
Altered pathways in OB derived from humans
 with COVID-19
(Reactome terms; transcriptomics)
Extracellular matrix organization
Collagen formation
Signaling by Receptor Tyrosine Kinases
Collagen biosynthesis and modifying enzymes
ECM proteoglycans
Non-integrin membrane-ECM interactions
Integrin cell surface interactions
Degradation of the extracellular matrix
Assembly of collagen fibrils and other multimeric structures
Collagen chain trimerization
Collagen degradation
Elastic fibre formation
MET promotes cell motility
MET activates PTK2 signaling
Nervous system development
Axon guidance
Crosslinking of collagen fibrils
Laminin interactions
Vesicle-mediated transport
Signaling by MET
MAPK family signaling cascades
EPH-Ephrin signaling
MAPK1/MAPK3 signaling
Neutrophil degranulation
Anchoring fibril formation
RAF/MAP kinase cascade
Regulation of cholesterol biosynthesis by SREBP (SREBF)
Signaling by NTRKs
Neuronal System
Signaling by NTRK1 (TRKA)
Activation of gene expression by SREBF (SREBP)
L1CAM interactions
GABA receptor activation
Membrane Trafficking
Immunoregulatory interactions between a Lymphoid and a non-Lymphoid cell
Asparagine N-linked glycosylation
Opioid Signalling
Toll-like Receptor Cascades
Diseases of signal transduction by growth factor receptors and second messengers
Integration of energy metabolism
Nuclear Events (kinase and transcription factor activation)
RET signaling
Aquaporin-mediated transport
Transport to the Golgi and subsequent modification
Negative regulation of MAPK pathway
Potassium Channels
Chaperonin-mediated protein folding
G-protein mediated events
Protein folding
CaM pathway
Calmodulin induced events
Protein-protein interactions at synapses
Ca-dependent events
PLC beta mediated events
DAG and IP3 signaling
Signaling by Nuclear Receptors
Neurotransmitter receptors and postsynaptic signal transmission
NGF-stimulated transcription
Transmission across Chemical Synapses
Intracellular signaling by second messengers
G alpha (s) signalling events
Signaling by WNT
TCF dependent signaling in response to WNT
EPH-ephrin mediated repulsion of cells
PIP3 activates AKT signaling
Interaction between L1 and Ankyrins
Signaling by FGFR
Supplementary figure 3

### Slide 4
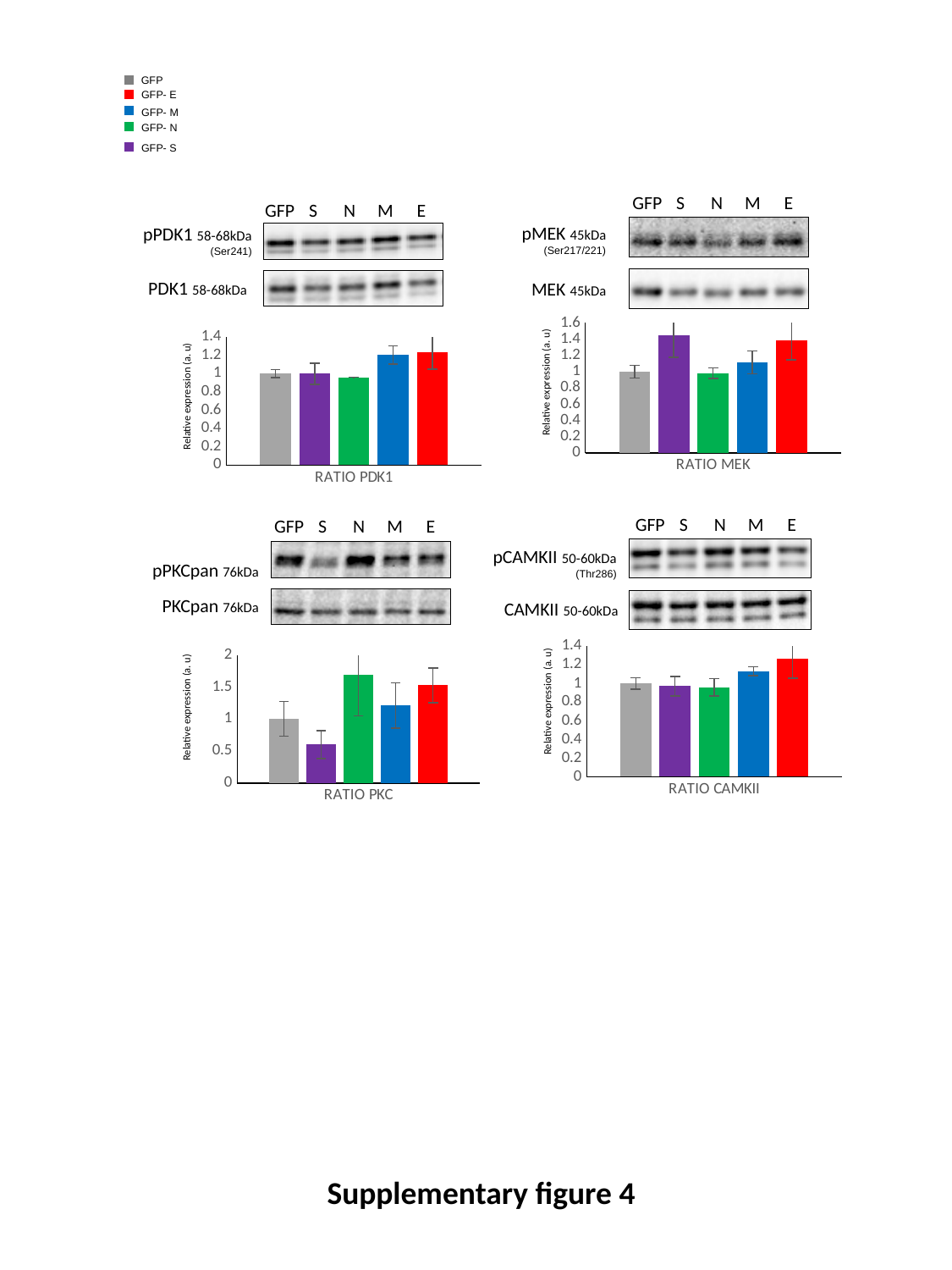

GFP
GFP- E
GFP- M
GFP- N
GFP- S
GFP
S
N
M
E
GFP
S
N
M
E
pMEK 45kDa
(Ser217/221)
pPDK1 58-68kDa
(Ser241)
PDK1 58-68kDa
MEK 45kDa
#### Chart
| Category | GFP | S | N | M | E |
|---|---|---|---|---|---|
| RATIO MEK | 1.0000000000000002 | 1.4503212863641775 | 0.983386909326753 | 1.1146713555797454 | 1.3845914803624815 |
#### Chart
| Category | GFP | S | N | M | E |
|---|---|---|---|---|---|
| RATIO PDK1 | 1.0000000000000002 | 0.9969597161265127 | 0.9595721547834222 | 1.2026972177745223 | 1.2299312367082929 |Relative expression (a. u)
Relative expression (a. u)
GFP
S
N
M
E
GFP
S
N
M
E
pCAMKII 50-60kDa
(Thr286)
pPKCpan 76kDa
PKCpan 76kDa
CAMKII 50-60kDa
#### Chart
| Category | GFP | S | N | M | E |
|---|---|---|---|---|---|
| RATIO CAMKII | 1.0 | 0.972242041194614 | 0.9600815217012081 | 1.1318049503301884 | 1.2635535026414628 |
#### Chart
| Category | GFP | S | N | M | E |
|---|---|---|---|---|---|
| RATIO PKC | 1.0 | 0.5996024413806171 | 1.6916265044978553 | 1.2094866282893346 | 1.5243999993960073 |Relative expression (a. u)
Relative expression (a. u)
Supplementary figure 4

### Slide 5
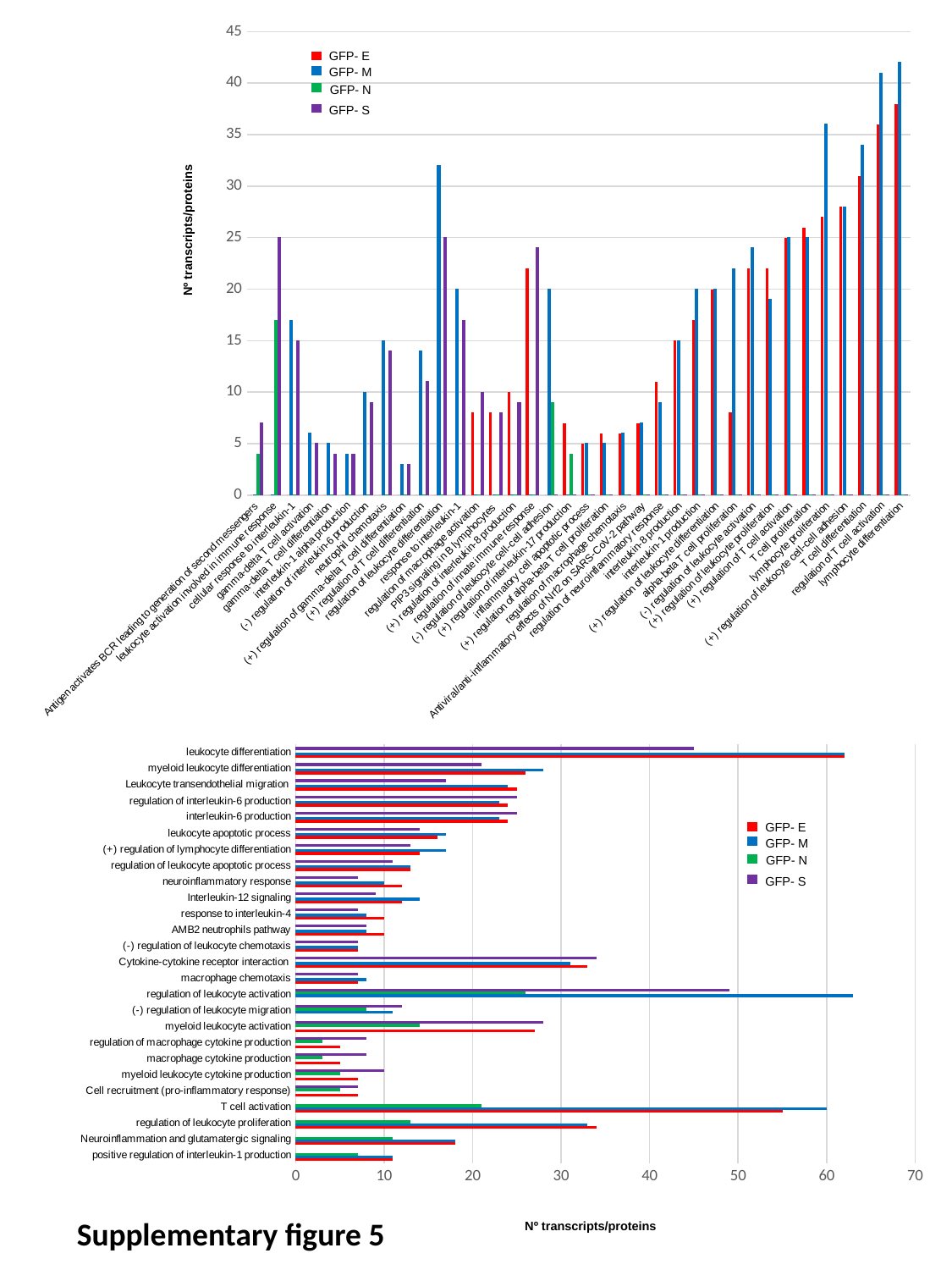

#### Chart
| Category | E PROTEIN | M PROTEIN | N PROTEIN | S PROTEIN |
|---|---|---|---|---|
| Antigen activates BCR leading to generation of second messengers | 0.0 | 0.0 | 4.0 | 7.0 |
| leukocyte activation involved in immune response | 0.0 | 0.0 | 17.0 | 25.0 |
| cellular response to interleukin-1 | 0.0 | 17.0 | 0.0 | 15.0 |
| gamma-delta T cell activation | 0.0 | 6.0 | 0.0 | 5.0 |
| gamma-delta T cell differentiation | 0.0 | 5.0 | 0.0 | 4.0 |
| interleukin-1 alpha production | 0.0 | 4.0 | 0.0 | 4.0 |
| (-) regulation of interleukin-6 production | 0.0 | 10.0 | 0.0 | 9.0 |
| neutrophil chemotaxis | 0.0 | 15.0 | 0.0 | 14.0 |
| (+) regulation of gamma-delta T cell differentiation | 0.0 | 3.0 | 0.0 | 3.0 |
| (+) regulation of T cell differentiation | 0.0 | 14.0 | 0.0 | 11.0 |
| regulation of leukocyte differentiation | 0.0 | 32.0 | 0.0 | 25.0 |
| response to interleukin-1 | 0.0 | 20.0 | 0.0 | 17.0 |
| regulation of macrophage activation | 8.0 | 0.0 | 0.0 | 10.0 |
| PIP3 signaling in B lymphocytes | 8.0 | 0.0 | 0.0 | 8.0 |
| (+) regulation of interleukin-8 production | 10.0 | 0.0 | 0.0 | 9.0 |
| regulation of innate immune response | 22.0 | 0.0 | 0.0 | 24.0 |
| (-) regulation of leukocyte cell-cell adhesion | 0.0 | 20.0 | 9.0 | 0.0 |
| (+) regulation of interleukin-17 production | 7.0 | 0.0 | 4.0 | 0.0 |
| inflammatory cell apoptotic process | 5.0 | 5.0 | 0.0 | 0.0 |
| (+) regulation of alpha-beta T cell proliferation | 6.0 | 5.0 | 0.0 | 0.0 |
| regulation of macrophage chemotaxis | 6.0 | 6.0 | 0.0 | 0.0 |
| Antiviral/anti-inflammatory effects of Nrf2 on SARS-CoV-2 pathway | 7.0 | 7.0 | 0.0 | 0.0 |
| regulation of neuroinflammatory response | 11.0 | 9.0 | 0.0 | 0.0 |
| interleukin-8 production | 15.0 | 15.0 | 0.0 | 0.0 |
| interleukin-1 production | 17.0 | 20.0 | 0.0 | 0.0 |
| (+) regulation of leukocyte differentiation | 20.0 | 20.0 | 0.0 | 0.0 |
| alpha-beta T cell proliferation | 8.0 | 22.0 | 0.0 | 0.0 |
| (-) regulation of leukocyte activation | 22.0 | 24.0 | 0.0 | 0.0 |
| (+) regulation of leukocyte proliferation | 22.0 | 19.0 | 0.0 | 0.0 |
| (+) regulation of T cell activation | 25.0 | 25.0 | 0.0 | 0.0 |
| T cell proliferation | 26.0 | 25.0 | 0.0 | 0.0 |
| lymphocyte proliferation | 27.0 | 36.0 | 0.0 | 0.0 |
| (+) regulation of leukocyte cell-cell adhesion | 28.0 | 28.0 | 0.0 | 0.0 |
| T cell differentiation | 31.0 | 34.0 | 0.0 | 0.0 |
| regulation of T cell activation | 36.0 | 41.0 | 0.0 | 0.0 |
| lymphocyte differentiation | 38.0 | 42.0 | 0.0 | 0.0 |GFP- E
GFP- M
GFP- N
GFP- S
Nº transcripts/proteins
#### Chart
| Category | E PROTEIN | M PROTEIN | N PROTEIN | S PROTEIN |
|---|---|---|---|---|
| positive regulation of interleukin-1 production | 11.0 | 11.0 | 7.0 | 0.0 |
| Neuroinflammation and glutamatergic signaling | 18.0 | 18.0 | 11.0 | 0.0 |
| regulation of leukocyte proliferation | 34.0 | 33.0 | 13.0 | 0.0 |
| T cell activation | 55.0 | 60.0 | 21.0 | 0.0 |
| Cell recruitment (pro-inflammatory response) | 7.0 | 0.0 | 5.0 | 7.0 |
| myeloid leukocyte cytokine production | 7.0 | 0.0 | 5.0 | 10.0 |
| macrophage cytokine production | 5.0 | 0.0 | 3.0 | 8.0 |
| regulation of macrophage cytokine production | 5.0 | 0.0 | 3.0 | 8.0 |
| myeloid leukocyte activation | 27.0 | 0.0 | 14.0 | 28.0 |
| (-) regulation of leukocyte migration | 0.0 | 11.0 | 8.0 | 12.0 |
| regulation of leukocyte activation | 0.0 | 63.0 | 26.0 | 49.0 |
| macrophage chemotaxis | 7.0 | 8.0 | 0.0 | 7.0 |
| Cytokine-cytokine receptor interaction  | 33.0 | 31.0 | 0.0 | 34.0 |
| (-) regulation of leukocyte chemotaxis | 7.0 | 7.0 | 0.0 | 7.0 |
| AMB2 neutrophils pathway | 10.0 | 8.0 | 0.0 | 8.0 |
| response to interleukin-4 | 10.0 | 8.0 | 0.0 | 7.0 |
| Interleukin-12 signaling | 12.0 | 14.0 | 0.0 | 9.0 |
| neuroinflammatory response | 12.0 | 10.0 | 0.0 | 7.0 |
| regulation of leukocyte apoptotic process | 13.0 | 13.0 | 0.0 | 11.0 |
| (+) regulation of lymphocyte differentiation | 14.0 | 17.0 | 0.0 | 13.0 |
| leukocyte apoptotic process | 16.0 | 17.0 | 0.0 | 14.0 |
| interleukin-6 production | 24.0 | 23.0 | 0.0 | 25.0 |
| regulation of interleukin-6 production | 24.0 | 23.0 | 0.0 | 25.0 |
| Leukocyte transendothelial migration  | 25.0 | 24.0 | 0.0 | 17.0 |
| myeloid leukocyte differentiation | 26.0 | 28.0 | 0.0 | 21.0 |
| leukocyte differentiation | 62.0 | 62.0 | 0.0 | 45.0 |GFP- E
GFP- M
GFP- N
GFP- S
Supplementary figure 5
Nº transcripts/proteins
